# A century of breeding has preserved genetic variation, accumulated favorable alleles, and shaped the *Rht* genes portfolio in North American spring wheat

**DOI:** 10.1101/2025.09.22.677111

**Authors:** Harsimardeep S. Gill, Sarah Blecha, Emily Conley, Charlotte Brault, Jason Fiedler, Jason Cook, Karl Glover, Andrew Green, Andrew Read, James A. Anderson

## Abstract

Hard red spring wheat (HRSW) is an important market class of wheat in North America with a rich history of breeding. We genotyped 1,013 HRSW lines representing a century of breeding material using a SNP array and targeted KASP genotyping to assess the changes in genetic diversity and *Rht* genes in HRSW from North America. Results show that early breeding efforts broadened an initially narrow gene pool derived from a few founders and subsequently maintained the genetic diversity. Analysis of *Rht* genes revealed that *Rht-D1b* was the predominant semi-dwarf allele from introduction of the Green Revolution genes (*Rht-B1* and *Rht-D1*) in 1960s until the FHB epidemics of 1990s, when most breeding programs shifted to *Rht-B1b*. However, adoption of *Rht-B1b* and *Rht-D1b* remains low in some regions of the Great Plains. The dwarf alleles at GA-sensitive *Rht24* and *Rht25* have persisted in the region at low frequencies before the introduction of Green Revolution genes and have increased in frequency in recent decades. We also analyzed the effect of *Rht* genes on plant height (PH) in the HRSW growing region and found that *Rht-B1b* or *Rht-D1b* results in ∼12% reduction in PH, which decreases to ∼9% in dry environments. Notably, combining dwarfing alleles at *Rht24* and *Rht25* reduced height by ∼6% in dry environments, suggesting an alternative approach for reducing PH. Our analysis also revealed significant genetic interactions between *Rht-B1* and *Rht25*, as well as between *Rht-D1* and *Rht24,* which could help fine-tune PH for different environments by employing appropriate *Rht* combinations. Furthermore, our results demonstrate positive selection for photoperiod sensitivity in recent decades, particularly in high-latitude regions. This study provides a valuable genomic resource, and the findings enhance our understanding of past breeding efforts while offering insights to guide future plant breeding strategies.

## 1. Introduction

Wheat (*Triticum aestivum* L.) is one of the world’s most important staple crops and plays a crucial role in global agriculture and food security (Shiferaw *et al*., 2013; Yao *et al*., 2025). Hard red spring wheat (HRSW) is the predominant market class of wheat in North America, known for its high grain protein content and strong gluten, making it ideal for bread baking. Historically, HRSW cultivation facilitated the agricultural expansion of the northern Great Plains with the introduction of the historic ‘Red Fife’ in 1842, which became the foundation of spring wheat production across the Northern U.S. in the late 19th century (Paulsen and Shroyer, 2008). Today, HRSW is a key contributor to regional and global food security, accounting for a significant share (∼30%) of U.S. wheat exports (USDA ERS, 2023). This importance highlights the need for ongoing genetic improvement in HRSW and encourages a better understanding of how a century of breeding has shaped its genetic structure and trait architecture.

Over the past century, wheat breeding has played a pivotal role in increasing global wheat productivity. A cornerstone of successful plant breeding programs lies in the ability to maintain and utilize genetic diversity, which allows breeders to respond effectively to environmental changes and emerging challenges (Tanksley and McCouch, 1997; Lehnert *et al*., 2022). However, intensive selection using a limited number of elite parents may inadvertently narrow the genetic base of a breeding program, potentially affecting long-term genetic progress (Fu *et al*., 2005; van de Wouw *et al*., 2010). For North American HRSW, the number of founder lines was relatively limited, with a few introductions from the 19th century making a disproportionate contribution to modern cultivars. As reported by Mercado et al (1996), three progenitors, namely ‘Red Fife’, ‘Hard Red Calcutta’, and ‘Turkey Red’, accounted for approximately 39% of the genetic lineage of HRSW. Therefore, it is essential to understand how genetic diversity has changed over a century of breeding efforts and whether plant breeding has had a negative impact on genetic variation in the HRSW region. It is also important to examine the genetic relatedness among different breeding programs within the HRSW region, as it offers insight into the regional overview of genetic variation. A few previous studies have shown mixed evidence regarding genetic erosion in the HRSW from North America (Fu *et al*., 2006; Sthapit *et al*., 2020, 2022; Semagn *et al*., 2021). However, none of these studies used a large set of HRSW germplasm that would reflect the genetic variation evolved through decades of breeding.

In addition to overall genetic variation, examining changes in allele frequency at specific genes that control important agronomic and developmental traits is crucial for gathering insights into the adaptation of HRSW in the Great Plains, the region that encompasses a wide range of climatic conditions. The most notable among these genes are the Reduced Height (*Rht*) genes, particularly the *Rht-B1b* and *Rht-D1b* semi-dwarf alleles introduced from the Japanese cultivar ‘Norin 10’, which became major drivers of the Green Revolution (Khush, 2001). These *Rht* genes encode defective DELLA growth proteins in the gibberellin (GA) hormone pathway, resulting in GA insensitivity, shorter plant stature, improved harvest index and increased grain yield under optimal conditions (Peng *et al*., 1999). Despite their global success, *Rht-B1b* and *Rht-D1b* have drawbacks, including reduced coleoptile length, emergence issues in dry soils, and potential yield penalties, particularly under stress conditions (Addisu *et al*., 2010; Würschum *et al*., 2017). The *Rht-B1b* and *Rht-D1b* were introduced into the HRSW region in the 1960s. However, no comprehensive study has investigated the distribution, deployment, and effects of these genes across varying geographical regions of the Northern Great Plains.

Another set of *Rht* genes, including *Rht24* and *Rht25*, which belong to the GA-sensitive group and reduce plant height without compromising stress tolerance, have been proposed as an alternative dwarfing system for dry environments (Würschum *et al*., 2017; Mo *et al*., 2018; Tian *et al*., 2022; Zhang *et al*., 2023). *Rht24* encodes a gibberellin 2-oxidase (TaGA2ox-A9), and its dwarfing allele (*Rht24b*) increases stem expression to lower bioactive GA levels and reduce plant height without affecting yield (Tian et al., 2022). *Rht25*, designated as PLATZ-A1 (TraesCS6A02G156600), is expressed mainly in the elongating stem and developing spike and encodes a plant-specific AT-rich sequence– and zinc-binding (PLATZ) protein, with its loss-of-function mutants conferring GA-sensitive dwarfing (Zhang *et al*., 2023). Recent studies have demonstrated the presence of dwarfing alleles of *Rht24* and *Rht25* in modern wheat, including various U.S wheat classes (Tian *et al*., 2019; Zhang *et al*., 2023). Moreover, Zhang *et al*. (2023) observed that two mutant alleles of *Rht25*, namely *Rht25b* and *Rht25f*, are prevalent in North American spring wheat. However, there is no information regarding the origin and distribution of alternative *Rht* genes in the HRSW from North America, nor their effect on plant height across regional environments in the presence or absence of *Rht-B1b* and *Rht-D1b*. Other genes, such as those involved in the vernalization response (*Vrn*) and photoperiod sensitivity (*Ppd*), play a crucial role in wheat’s adaptation to diverse environments. Among the group of *Vrn* and *Ppd* genes, allelic variations at homeologues of *VRN1* (*Vrn-A1*, *Vrn-B1*, *Vrn-D1*) and *PPD1* are the main contributors to the flowering time and acclimatization of wheat to specific environments (Trevaskis, 2010; Grogan *et al*., 2016). Fine-tuning the different combinations of *Vrn* and *Ppd* genes reflects breeding priorities, and tracking their allele frequency is critical for evaluating breeding impacts and guiding future strategies.

To address these research gaps, an extensive material and historical dataset for HRSW is needed. The Hard Red Spring Wheat Uniform Regional Nursery (HRSWURN) offers a valuable resource, capturing nearly a century of coordinated wheat breeding in North America. Initiated in the late 1920s, the HRSWURN annually evaluates elite breeding lines representing major public and private breeding programs across the northern Great Plains. This nursery encompasses not only commercial cultivar releases but also advanced breeding lines, offering a valuable resource for calculating accurate genetic diversity estimates over several decades. Despite the availability of extensive phenotypic data from this nursery (Yao *et al*., 2022; Gill *et al*., 2025), genotypic characterization and genotype-informed analysis of this historical germplasm collection remain unexplored. In this study, we collected and genotyped 1,013 HRSW accessions evaluated in the HRSWURN over the past century, representing material developed and tested from 1912 to 2023. We generated genome-wide genotype data using a low-density 3K SNP array, complemented with targeted KASP assays to characterize important functional genes. Specifically, we aimed to (i) elucidate genetic relatedness and population structure among major spring wheat breeding programs in Northern America, (ii) assess genome-wide genetic diversity and quantify its temporal dynamics over the past century, (iii) investigate shifts in allelic frequencies of key genes related to developmental traits, and (iv) study the origin and distribution of various *Rht* genes in this region and evaluate their effect on plant height across diverse environments of the Northern Great Plains.

## 2. Materials and Methods

### 2.1 Description of the historical HRSWURN material

The HRSWURN, coordinated by the United States Department of Agriculture – Agricultural Research Service (USDA-ARS) researchers, allows various public and private breeding programs to contribute their advanced material for evaluation in multi-environment trials. The USDA-ARS compiles the multilocation data and publishes it in a comprehensive annual report, which is distributed to contributing breeders and made publicly available on the GrainGenes website (https://graingenes.org/GG3/).

In this study, we collected a large set of accessions that have been evaluated in HRSWURN trials from 1929 to 2023. Although HRSWURN testing officially began in 1929, several earlier-released lines, such as ‘Marquis’ from 1912, were included as checks in the nursery’s initial years. The seed of historical HRSWURN entries was sourced from public breeding programs and the National Plant Germplasm System. A total of 1,013 accessions were collected and genotyped (Supplementary Table S1). These included material from four major public HRSW breeding programs in the U.S., including those of North Dakota (ND, nL=L355), Minnesota (MN, nL=L219), South Dakota (SD, nL=L157), and Montana (MT, nL=L85). Additionally, 61 lines from Canadian public programs (AAFC), 123 lines from thirteen different seed companies (PRIVATE), and 13 lines from small programs such as Wisconsin were included (Figure 1). We assigned each accession to a decade bin (e.g., “1930s” = 1930-1939) based on its first year of inclusion in the HRSWURN. In total, the collection averages approximately 12 lines per year and 101 lines per decade, and this panel captures the temporal breadth and program-wide diversity of HRSW breeding over nearly a century.

**Figure 1.**
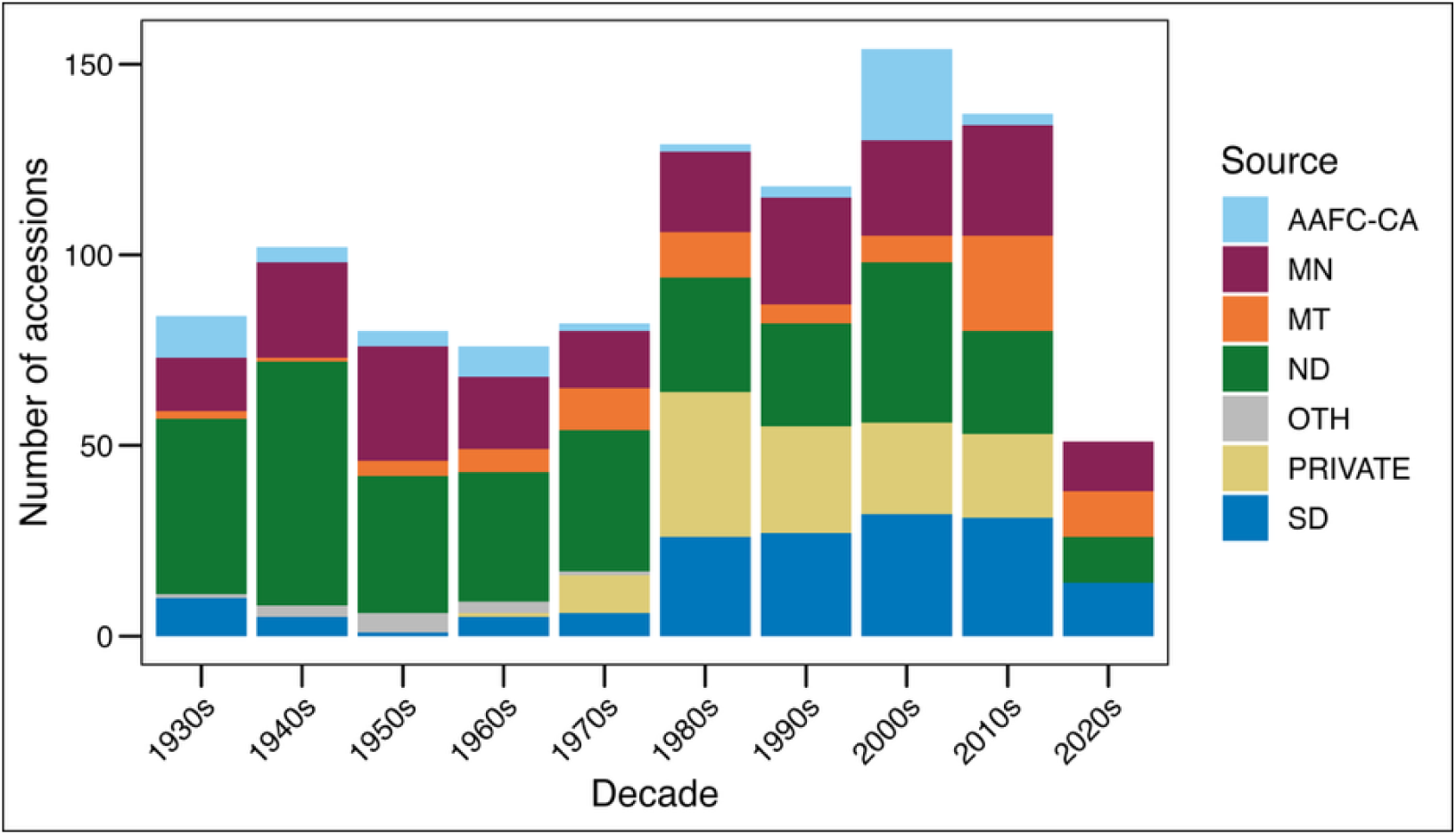
Sources of test entries in the Hard Red Spring Wheat Uniform Regional Nursery (HRSWURN) from the 1930s to the 2020s that were collected and genotyped in this study. AAFC-CA, Canadian public breeding programs; MN, Minnesota; MT, Montana; ND, North Dakota; SD, South Dakota; OTH, Other public breeding programs, including Idaho, Wisconsin, and a few CIMMYT entries; PRIVATE, the lines contributed by private sector seed companies.

### 2.2 Genotyping analysis

Leaf tissue was collected from six seedlings per accession at the twolZI to threelZIleaf stage and bulked for DNA extraction. Genotyping was performed on the USDAlZISoyWheOatBarlZI3K Infinium® II array (Illumina), which contains ∼3,421 evenly spaced, highlZIinformation markers across the wheat genomes. Single-nucleotide polymorphisms (SNPs) were called using the Genome Studio Polyploid Clustering Module v1.0 (Illumina 2013), after manual curation to ensure that cluster positions matched the fluorescence data. Marker positions were assigned based on the Basic Local Alignment Search Tool (BLAST) (Altschul *et al*., 1990) against the IWGSC RefSeq v2.1 reference genome (Zhu *et al*., 2021). In total, 3,107 raw SNP calls were generated across 21 wheat chromosomes. The SNPs were subjected to quality control using TASSEL v5.0 (Bradbury *et al*., 2007). First, the markers were filtered for more than 30% missing data and more than 25% heterozygosity; however, there were no markers to exclude based on these criteria. Second, 606 markers were removed based on a minor allele frequency (MAF) of less than 1%, resulting in 2,501 markers for further analysis. These 2,501 markers were not imputed for genetic relatedness and diversity analysis to avoid any bias in estimates due to imputation.

Additionally, the panel was genotyped for genes associated with various agronomic traits. This included different *Rht* genes, such as *Rht-B1*, *Rht-D1*, *Rht*8, *Rht24*, and *Rht25*. For *Rht25*, we focused on two different natural variants, namely *Rht25b* and *Rht25f*, based on their higher frequencies in spring wheat (Zhang *et al.,* 2023). The panel was also screened for allelic variation at genes related to photoperiod sensitivity (*PPD1*), vernalization response (*VRN1*), and genes associated with flowering time (*Vrn-A3, Vrn-D3, TaELF3-B1, TaELF3-D1,* and *TaFT3-B1*). Due to negligible variation, we excluded the *Vrn-A3, TaELF3-B1,* and *TaELF3-D1* from the results section. The functional markers for some of these genes are included in the 3K array; however, most were genotyped using Kompetitive Allele-Specific PCR (KASP) platform using published oligos. A detailed summary of the functional markers used in the study, along with their references, is provided in Supplementary Tables S2 (markers included in the 3K array) and S3 (KASP-based markers).

### 2.3 Population structure and genetic relatedness analysis

Principal component analysis (PCA) was conducted with the SNPRelate package <u>(Zheng et al. 2012)</u> in R version 4.4.0. We used the first two principal components to visualize genetic relationships among accessions across decades and breeding programs (R Core Team, 2023). In addition, Bayesian clustering was performed using STRUCTURE v2.3.4 (Pritchard *et al*., 2000), assuming up to 10 subpopulations (K = 1–10). For each K, ten independent replicates were performed, comprising a burnlZIin of 20,000 iterations followed by 20,000 MCMC iterations. The most likely number of clusters was determined using the ΔK method through the offline implementation of the STRUCTURE Harvester <u>(Earl and vonHoldt 2012)</u>. Furthermore, we conducted an analysis of molecular variance (AMOVA) to quantify genetic partitioning among groups defined by both decade and breeding program. AMOVA was performed in R, utilizing the ‘poppr’ package with 999 permutations (Kamvar *et al*., 2014). Pairwise genetic differentiation (*F_ST_*) among the major breeding programs was estimated using the Weir and Cockerham method using the R package ‘hierfstat’ (Weir and Cockerham, 1984; Goudet and Jombart, 2004).

### 2.4 Estimation of genetic diversity

For estimation of genetic diversity, we followed the approach documented by Sthapit et al. (2020).

Genetic diversity within each decade was assessed by calculating expected heterozygosity (*H_e_*; equivalent to gene diversity) at the genome level. For a locus with n alleles, *H_e_*is defined as

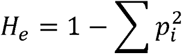

where *H_e_* is the gene diversity at a locus and *p_i_* is the frequency of the *i*^th^ allele in the population. As explained by Sthapit et al. (2020), gene diversity is a suitable measure of genetic diversity for self-pollinated crops and closely related populations (Nei, 1973; Weir, 1996). Gene diversity is mathematically equivalent to the polymorphism information content (PIC) in the case of a biparental population. We computed gene diversity per decade using the hierfstat package in R. The temporal trends in genetic diversity were evaluated by grouping the accessions by decade, as described in the previous section.

### 2.5 Statistical analysis

#### 2.5.1 Phenotypic analysis of historical plant height data

The detailed description of the experimental setup for the historical HRSWURN trials can be found in Gill et al. (2025). We utilized historical plant height data from the HRSWURN trials, spanning from 1968 to 2023, to assess the impact of *Rht* alleles on plant height in the HRSW region. We used data from 1968, as the nursery included two long-term checks (‘Chris’ and Marquis) from 1968 to 2023, along with several short-term checks, which allowed for the estimation of year effects and obtaining genotype means over the years. First, we utilized the available data from all tested locations to estimate the plant height best linear unbiased estimates (BLUEs) for 768 unique genotypes tested from 1968 to 2023, for which genotype data were available. For this, a linear mixed model was used as described in the following equation:

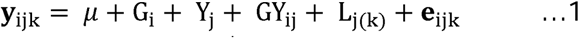

where y_ijk_ is adjusted entry mean of the i^th^ genotype in year j at site k obtained from the first stage of analysis, *µ* is the overall mean, G_i_ is the effect of the i^th^ genotype, Y_j_ is the effect of the j^th^ year, GY_ij_ is the genotype-by-year interaction effect, L_jk_ is the effect of location k nested within year j, and e_ijk_ is the residual, due to the combined effects of within-trial error and genotype × location within-year interaction. Both genotype and year effects were considered fixed, as treating them as random effects could introduce bias. As suggested by Mackay et al. (2011), treating these as random can lead to biased variance estimates and affect the distribution of variety effects due to anticipated large and potentially non-linear trends over time for both genotype and year effects.

Additionally, we estimated the effect of *Rht* alleles at four different locations: Crookston (MN), Carrington (ND), Saint Paul (MN), and Williston (ND). These locations were chosen to represent the diverse moisture regimes across the Northern Great Plains region. The annual precipitation for each location from 1968 to 2023 was extracted using the ‘rnoaa’ package in R (https://github.com/ropensci/rnoaa), and the geographical information for these sites is presented in Supplementary Table S4. The plant height BLUEs for each of these four locations were derived using a mixed model that included genotype and year as fixed effects, with the genotype × year interaction as a random effect:

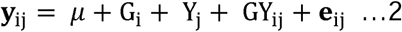

where y_ij_ is adjusted entry mean of the i genotype in year j obtained from the first stage of analysis (individual locations), *µ* is the overall mean, G_i_ is the effect of the i^th^ genotype, Y_j_ is the effect of the j^th^ year, GY_ij_ is the genotype-by-year interaction effect, and e_ij_ is the residual effect.

#### 2.5.2 Linear models for estimation of *Rht* allele effects

We evaluated the effects of dwarf alleles at *Rht*lZI*B1*, *Rht*lZI*D1*, *Rht24*, and *Rht25* loci on plant height following <u>Grogan et al. (2016)</u>. We evaluated the effect of *Rht* alleles using BLUEs derived from a combined multilZIenvironment analysis of HRSWURN (equation 1) and for the four individual locations (equation 2). Only accessions with complete, homozygous genotype calls at all four *Rht* loci were included. A global linear model, including the main effects of all four genes and all possible two-way interactions, was fit using plant height BLUEs as the response variable with the ‘lm’ function in R. The global model was subjected to model selection using the ‘dredge’ function in the MuMIn package (Bartoń, 2010) to determine the best fit, as indicated by the corrected Akaike Information Criterion (AICc). Furthermore, the best model was fitted using BLUEs derived from the combined analysis, and for BLUEs derived from Crookston (CRK), Carrington (CRT), Saint Paul (STP), and Williston (WLT), to estimate the intercept and quantify the effect of *Rht* alleles. In some cases, the selected best model included non-significant terms; however, removing these terms detracted from the model fit.

### 2.6 Genome**-**wide association using historical plant height data

A genome-wide association study (GWAS) was performed using a filtered set of 3,000 SNPs and plant height BLUEs obtained from Equation 1. The 2,501 markers (described in section 2.2) were further filtered for 5% minimum allele frequency and subjected to imputation using the LD-kNN method in TASSEL (Bradbury *et al*., 2007). A filtered and imputed set of 2,275 markers was used for GWAS with historical plant height data from 677 historical genotypes. We used a multilocus mixed model, Fixed and random model Circulating Probability Unification (FarmCPU), to perform GWAS (Liu *et al*., 2016; Wang and Zhang, 2021). The FarmCPU model was implemented using the Genomic Association and Prediction Integrated Tool (GAPIT) version 3.0 (Wang and Zhang, 2021) in the R environment, including the first two principal components to account for population structure. Marker-trait associations were considered significant based on a strict Bonferroni-corrected threshold of *P < 0.01* (–log_10_P = 5.36).

### 2.7 Software and code

All the statistical analyses were performed in R version 4.4.0 using the base or other relevant packages. The mixed model analysis was performed using ASREML-R version 4.2 (Butler *et al*., 2018). The linear model for analyzing the effect of *Rht* alleles was performed using the ‘lm’ function of R. The visualizations for allelic frequency changes were created using the R package ‘ggplot2’ (Wickham, 2016).

## 3. Results

### 3.1 Population structure and genetic relationships

Genotype data from the 3K array was subjected to quality control, resulting in 2,501 SNPs evenly distributed across the 21 wheat chromosomes (Supplementary Table S5). The A and B sub-genomes had 1,072 and 1,058 SNPs, respectively, whereas the D genome had a comparatively low representation with 371 SNPs (Supplementary Figure S1).

Principal component analysis (PCA) was performed using the biallelic SNP markers from the 3K array. The first two principal components from the genotypic data accounted for approximately 7% and 4.4% of the total variation (Figure 2A). PCA revealed a modest structure, highlighted by the year of origin, with Pre-1970 lines tending to cluster together and some overlap between the Pre– and Post-1970 groups (circles vs. triangles in Figure 2A). Visualization of the PCA biplot by decade, spanning from the 1930s to the 2020s, explained this overlap and revealed a more continuous temporal gradient along the principal components, rather than discrete clusters (Figure 2A), suggesting gradual rather than sudden shifts in allele frequencies.

**Figure 2.**
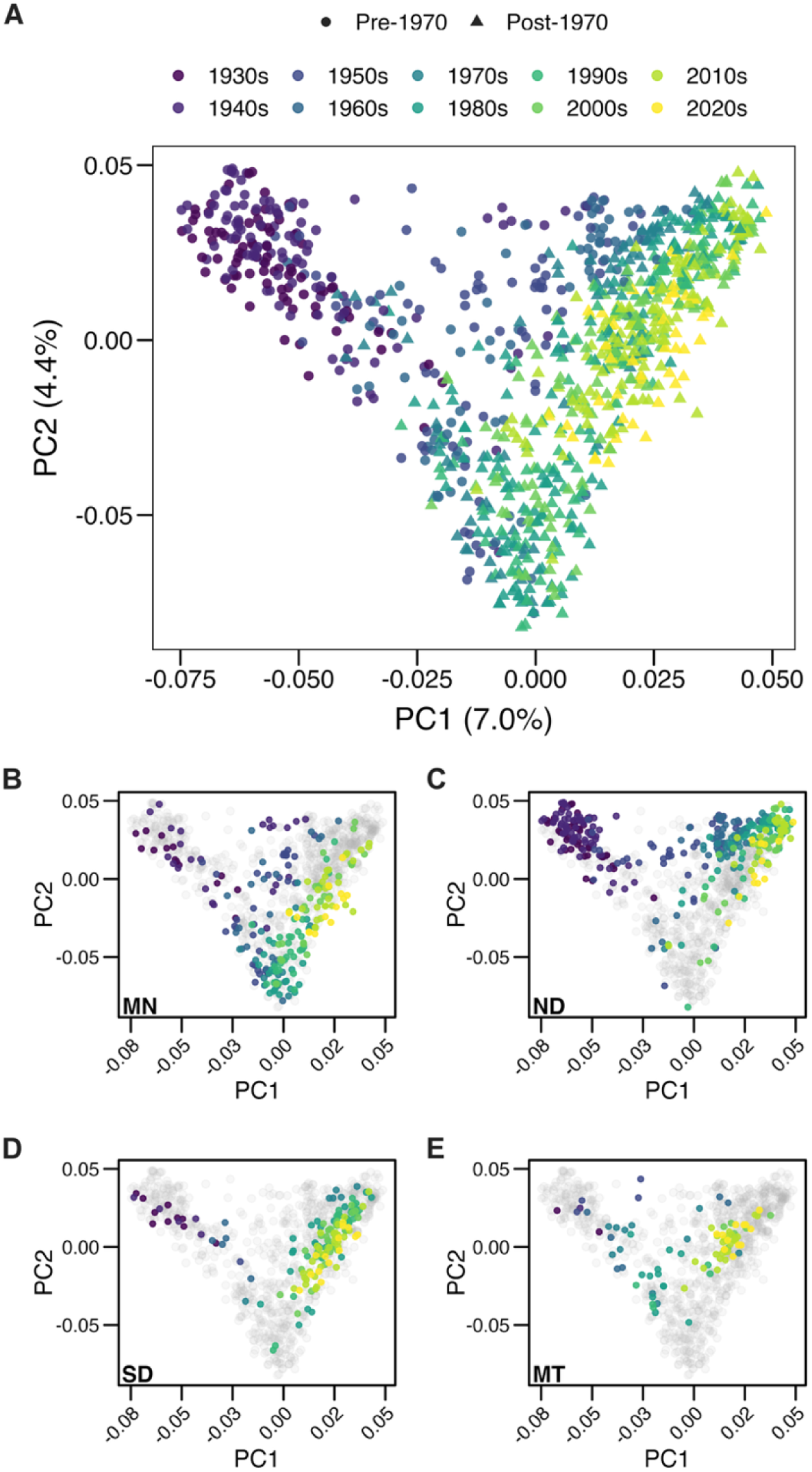
Principal component analysis of 1,013 accessions. (A) Principal component analysis (PCA) of spring wheat breeding germplasm showing the temporal differentiation. The PCA was conducted based on 3K SNP data and the variance explained by PC1 and PC2 is indicated in parentheses. Germplasm is categorized broadly as pre-1970 (circles) and post-1970 (triangles) and further subdivided by individual decades from the 1930s to the 2020s, with color gradients ranging from dark purple (older decades) to bright yellow (recent decades). (B-E) The genetic structure of spring wheat germplasm is illustrated by highlighting lines developed by four major public breeding programs of the region including Minnesota (MN), North Dakota (ND), South Dakota (SD), and Montana (MT) (colored by the decade of origin), against the overall germplasm from the HRSW region (grey dots).

We also used PCA to study the relatedness among MN, ND, SD, and MT breeding programs. The MN lines showed the broadest dispersion across both PCs, reflecting the historically diverse parentage employed in that program (Figure 2B). Notably, a compact cluster of related accessions in the 1980s–1990s was observed in the MN program, which seems very distinct from other regional breeding pools. In contrast, the ND program formed two closely clustered groups, one in the Pre-1970 era and the other in the Post-1970 era (Figure 2C). The SD and MT programs exhibited intermediate patterns and showed moderate clustering, with the SD program displaying a more scattered distribution (Figure 3). Additionally, the PCA revealed that materials from the four programs tend to cluster together in recent decades, with accessions from the 1990s, 2000s, and 2010s (green to yellow hues) projecting into relatively restricted sectors of the individual biplots (Figures 2B-2E). Furthermore, the STRUCTURE analysis suggested two major groups based on the ΔK statistic, which primarily separated the Pre-1970 and Post-1970 categories (Supplementary Figure S2). The results of PCA and STRUCTURE showed that although the temporal classes (Pre-1970 vs. Post-1970) correlate with the two major clusters, other factors also contribute to the observed genetic structure.

**Figure 3.**
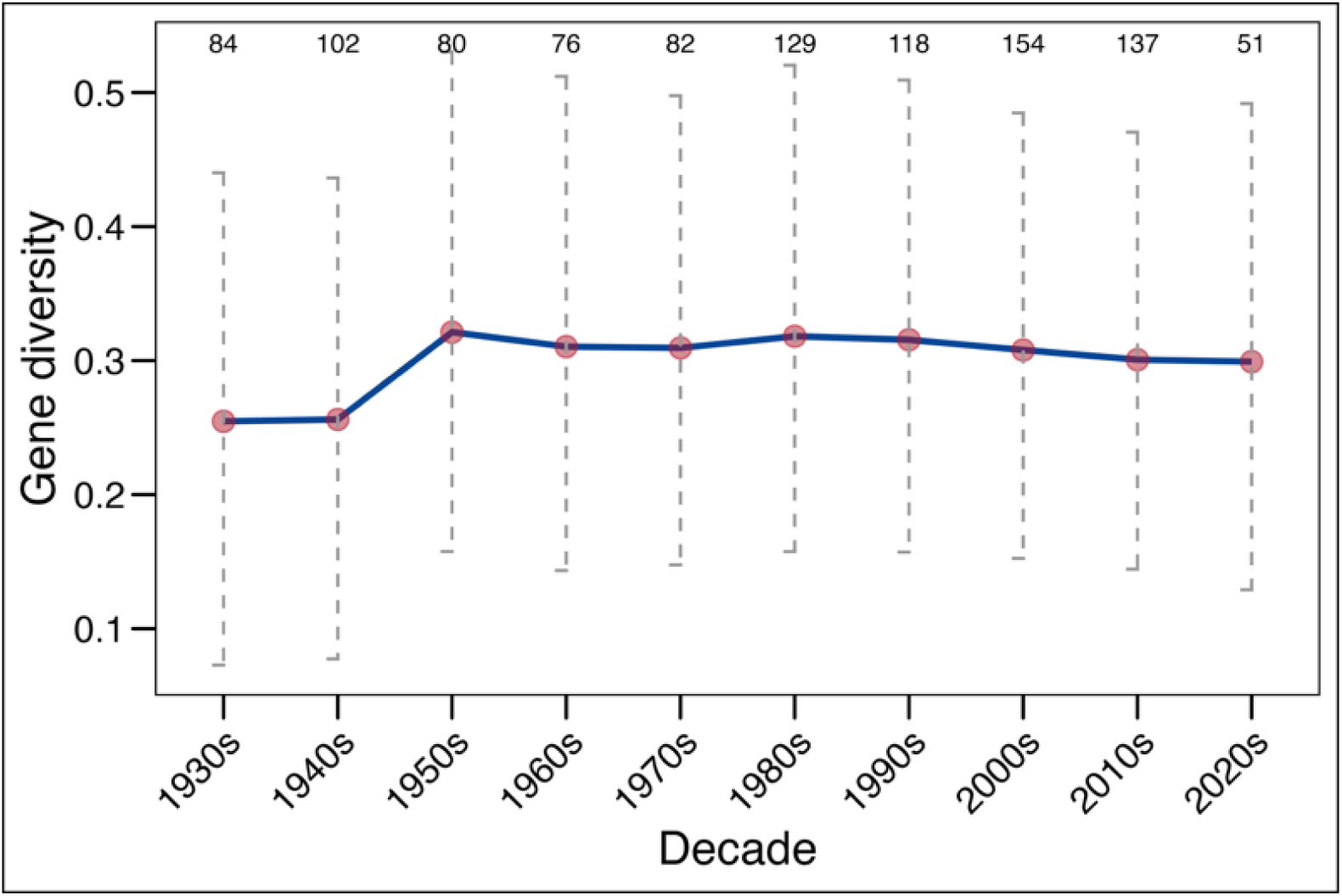
Overall genetic diversity (based on expected heterozygosity) in HRSW breeding germplasm evaluated across decades from the 1930s to the 2020s. The solid blue line represents the expected heterozygosity trend, with red points representing the mean and vertical dashed lines representing the standard deviation. The number at the top of each group represents the number of genotypes for the given decade.

### 3.2 Analysis of molecular variance (AMOVA) and genetic differentiation (*F_ST_*)

We conducted AMOVA by grouping the accessions based on temporal origin (Pre-1970 and Post-1970) and breeding program (Table 1). AMOVA for temporal classes revealed that lines released before and after 1970 are moderately differentiated, with the two classes accounting for 11% of the total variation (Phi = 0.11, p = 0.001). When lines were grouped by breeding programs from the entire region (MN, MT, ND, SD, AAFC-CA, and PRIVATE), only 6.3% of the total variation was attributed to the different breeding programs (Phi = 0.06, p = 0.001). In comparison, 93.7% of the variation was attributed within programs.

**Table 1.** Analysis of molecular variance (AMOVA) for the 1,013 accessions based on two groups, including two temporal classes (pre-1970 vs. post-1970 entries) and six breeding programs (MN, ND, SD, MT, AAFC-CA, and PRIVATE). For each group, the total variation is decomposed into a between-groups component and a within-groups component.

| Group | Source | Df | Sum Sq | Mean Sq | Sigma | % | Phi | p-value |
| --- | --- | --- | --- | --- | --- | --- | --- | --- |
| Pre-1970<br>v/s Post-1970 | Between<br>samples | 1 | 38418.7 | 38418.7 | 83.3 | 11.1 | 0.11 | 0.001 |
|  | Within<br>samples | 1011 | 674895.1 | 667.6 | 667.6 | 88.9 |  |  |
|  | Total | 1012 | 713313.8 | 704.9 | 750.9 | 100 |  |  |
| Breeding<br>programs | Between<br>samples | 5 | 38308.7 | 7661.7 | 45.1 | 6.3 | 0.06 | 0.001 |
|  | Within<br>samples | 994 | 662920.2 | 666.9 | 666.9 | 93.7 |  |  |
|  | Total | 999 | 701228.8 | 701.9 | 712.1 | 100 |  |  |

Df = degrees of freedom; Sum Sq = sum of squares; Mean Sq = mean squares; Sigma = estimated variance component; % = percentage of total molecular variance explained by that component; Phi ( $\Phi$ ) = $\Phi$ -statistic measuring the proportion of total genetic variance that lies among groups; p-value derived from 999 permutations of individuals among groups. A significant $\Phi$ ( $p < 0.05$ ) indicates departure from panmixia and the presence of genetic structure attributable to the grouping factor.

Next, we employed pairwise *F_ST_* to gain a better understanding of the genetic relatedness among breeding programs in the HRSW region. Pairwise *F_ST_* values among the various breeding programs ranged from 0.02 to 0.10 (Table 2). The results indicated varying degrees of genetic divergence among the programs, with the MN program exhibiting low genetic differentiation with all other programs, including the PRIVATE germplasm. The germplasm from MT showed a high genetic differentiation from all other public breeding programs, as well as PRIVATE germplasm. The PRIVATE germplasm demonstrated moderate differentiation with most public breeding programs, except for the MN program (*F_ST_* = 0.029). Interestingly, the AAFC-CA germplasm exhibited moderate differentiation from the U.S. public breeding programs and PRIVATE germplasm (Table 2).

**Table 2.** Pair-wise genetic differentiation *F_ST_* among different hard-red spring wheat breeding from North America. The table shows Weir & Cockerham’s unbiased *F_ST_*estimates calculated from genome-wide SNPs for each pair of breeding programs. Lower-triangle values represent the proportion of total genetic variance attributable to population structure between the two programs, and the upper triangle is left blank for clarity. An *F_ST_* of 0 indicates no detectable differentiation, whereas values ≥ 0.05, 0.10, and 0.25 are commonly interpreted as low, moderate, and high differentiation, respectively (Wright 1978).

| **Program** | **MN** | **ND** | **SD** | **MT** | **AAFC-CA** |
| --- | --- | --- | --- | --- | --- |
| ND | 0.050 | - | - | - | - |
| SD | 0.060 | 0.050 | - | - | - |
| MT | 0.070 | 0.060 | 0.100 | - | - |
| AAFC-CA | 0.050 | 0.050 | 0.080 | 0.090 | - |
| PRIVATE | 0.020 | 0.090 | 0.080 | 0.090 | 0.080 |

AAFC-CA, Canadian public breeding programs; MN, Minnesota; MT, Montana; ND, North Dakota; SD, South Dakota; and PRIVATE refers to the lines contributed by different seed companies from the private sector.

### 3.3 Temporal changes in genetic diversity

Genetic Diversity was inferred using SNP markers by calculating expected heterozygosity (*He)*. The temporal analysis across the entire HRSW region showed a significant increase in mean expected heterozygosity during the 1950s, followed by a periodic increase over the last seven decades (Figure 3, Supplementary Table S6). The *He* increased from 0.25 in the 1930s to a peak of 0.32 in the 1950s (Figure 3; Supplementary Table S6). It remained relatively high throughout the following decades, with mean values ranging from 0.31 to 0.32 from the 1960s to 2000s, followed by a slight decrease in recent decades (0.30). Despite these temporal fluctuations, our results demonstrate that plant breeding efforts have helped increase genetic diversity in HRSW germplasm from North America, which has been maintained over time.

We examined the genetic diversity in four major public breeding programs from MN, MT, ND, and SD. Importantly, we disregarded the lower estimates for the 2020s in these comparisons, as they are based on fewer accessions over only three years and may not be comparable to those from other decades. In the MN breeding program, mean expected heterozygosity increased substantially from 0.26 in the 1930s to a peak of 0.32 in the 1950s. Although it showed a decline in the following decades and reached a low of 0.26 in the 1980s, it exhibited a gradual increase in recent decades (Supplementary Table S7). The MT program had a mean expected heterozygosity of 0.19 in the 1930s, which peaked at 0.32 during the 1960s and 1990s, but there has been a significant decline in recent decades, with estimates dropping below 0.30. The ND program had maintained the mean expected heterozygosity values around 0.25 from the 1930s to the 2000s but experienced a decrease in the 2010s. The expected heterozygosity for SD program was 0.25 in the 1930s and 1940s, and the levels remained in the 0.27–0.29 range in more recent decades (Supplementary Table S7).

### 3.4 Temporal shifts in Reduced height (*Rht*) genes

#### 3.4.1 Distribution and frequency of *Rht* alleles in North American HRSW

The historical panel was genotyped for four *Rht* genes, *Rht-B1*, *Rht-D1*, *Rht24*, and *Rht25,* to analyze the frequency of height-reducing alleles over time, based on 972 lines with complete genotype calls (Supplementary Table S8). A subset of 340 accessions was also genotyped for *Rht8*, which revealed only two accessions with dwarf alleles and 338 lines with the tall allele. Due to negligible variation, *Rht8* was dropped from further analysis. The GA-insensitive dwarf alleles at the *Rht-B1* and *Rht-D1* loci (*Rht-B1b* and *Rht-D1b*) were introduced during the Green Revolution in the 1960s, with *Rht-D1b* emerging at a slightly higher frequency than *Rht-B1b* (Figures 4A and 4B). The dwarf allele at the *Rht-D1* locus (*Rht-D1b*) was favorably adopted in subsequent decades, with its frequency increasing dramatically and reaching ∼46% during the 1980s compared to only ∼23% for *Rht-B1b* (Figures 4A and 4B; Supplementary Table S8). However, the trend reversed in the 1990s with a continuous decline in the frequency of *Rht-D1b* and an increase in *Rht-B1b*. In the most recent two decades, *Rht-B1b* was present in more than 65% of HRSW germplasm, while the frequency of *Rht-D1b* during this time remained below 10% (Supplementary Table S8).

**Figure 4.**
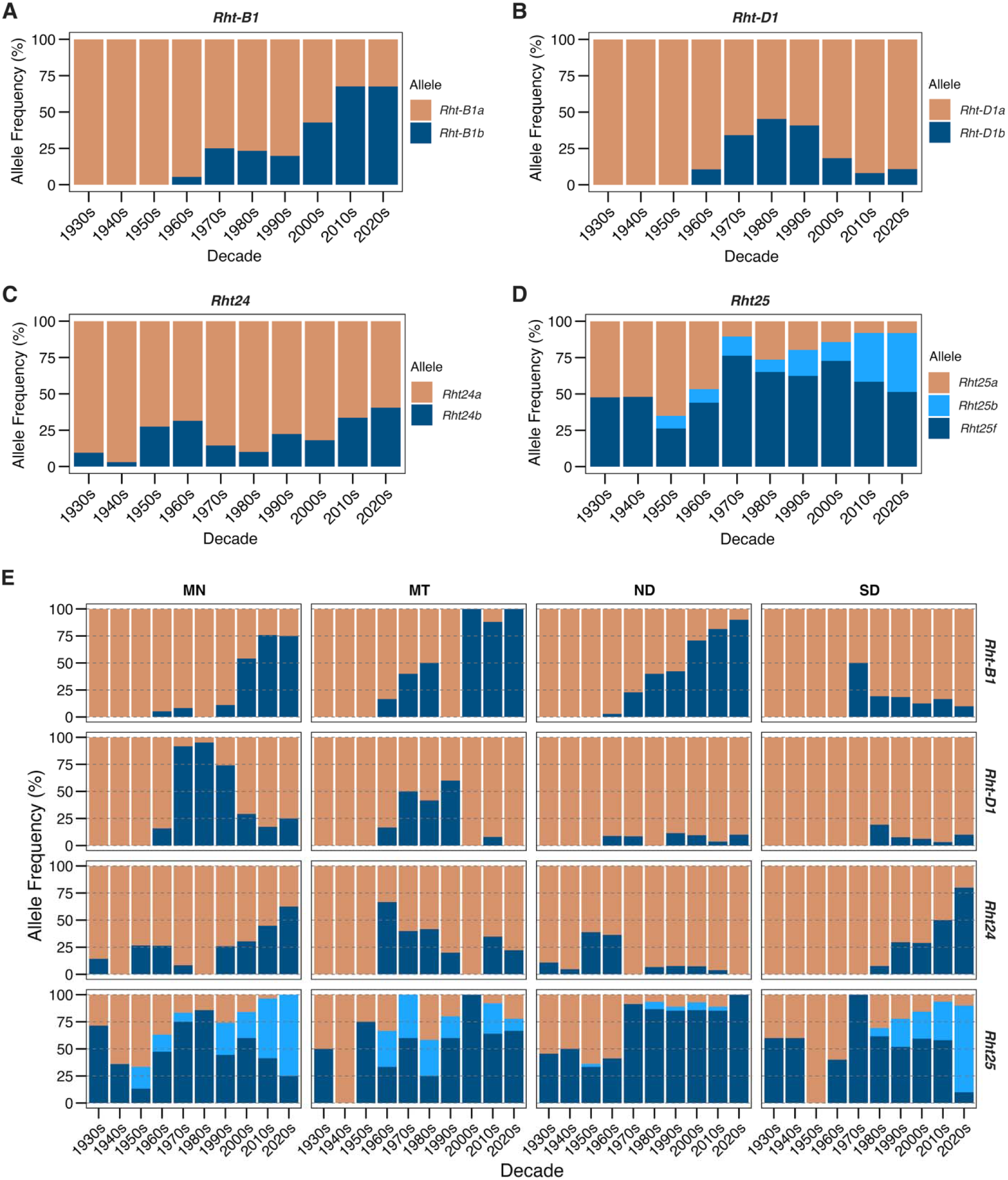
Temporal trends in allele frequency for four major reduced-height (*Rht*) genes in HRS wheat over the century. Panels show the overall allele frequency of (A) *Rht-B1*, (B) *Rht-D1*, (C) *Rht24*, and (D) *Rht25*. Panel E illustrates the changes in allelic frequency of these genes among four major public breeding programs of the region, including Minnesota, MN; Montana, MT; North Dakota, ND; and South Dakota, SD. Colors indicate different alleles for respective genes, with the orange color representing the wildtype/tall allele.

The dwarf allele at the GA-sensitive *Rht24* locus (*Rht24b*) has been present in the regional germplasm at low frequencies since the 1930s, well before the introduction of *Rht-B1b* and *Rht-D1b* (Figures 4A-4D). The frequency of *Rht24b* gradually increased until the 1960s but declined following the adoption of *Rht-B1b* and *Rht-D1b*. The frequency of *Rht24b* ranged from ∼10 to ∼20% in the region from the 1970s to the 2000s, until a sudden increase in the last 20 years when it reached up to 40% (Figure 4C). Similar to *Rht24*, one dwarf allele at the *Rht25* locus (*Rht25f*) has been present at high frequencies since the 1930s, with its frequency reaching approximately 75% in the 1970s. Interestingly, another dwarf allele at this locus (*Rht25b*), which first emerged in the 1950s, has consistently increased in frequency and replaced *Rht25f* in recent decades (Figure 4D).

Further analysis of *Rht* allele frequencies within each major breeding program revealed distinct adoption trajectories for the dwarfing alleles (Figure 4E, Supplementary Table S9). In the MN program, the frequency of *Rht-D1b* increased rapidly after its introduction in the 1960s, surpassing 90% by the 1980s. However, *Rht-D1b* declined to approximately 25% by the 2020s, as *Rht-B1b* was favored in recent decades, reaching around 75%. Despite its early introduction, *Rht24b* remained rare (<30%) until the 1990s; however, it has gradually increased over the past two decades. For *Rht25*, the dwarf allele *Rht25f* showed a consistent increase from the 1950s to the 1980s. However, another dwarf allele at this locus, *Rht25b*, has been highly favored in recent years and was present in over 70% of lines from MN during the past decade. In contrast to the MN program, the ND program adopted *Rht-B1b* instead of *Rht-D1b*, with the frequency of *Rht-B1b* steadily increasing to approximately 90% by the 2020s, while *Rht-D1b* remained at a very low frequency. The frequency of *Rht24b* has remained low, while *Rht25f* has been an important component of the ND program since the 1970s, reaching fixation by the 2020s.

The MT program deployed both *Rht-B1b* and *Rht-D1b* at nearly equal frequencies until the 1970s–1980s, after which *Rht-B1b* became highly favored and was almost fixed by the 2000s. The dwarf alleles at *Rht25*, especially *Rht25f*, remained dominant since the 1970s, still accounting for ∼67% in the 2020s. We observed an interesting pattern for the SD program regarding the adoption of green revolution genes (Figure 4E). With high early adoption, *Rht-B1b* was present in ∼50% of SD germplasm by 1980. However, the frequency of *Rht-B1b* and *Rht-D1b* did not exceed 20% after the 1980s. Conversely, the SD program has relied heavily on the combination of *Rht24b* and *Rht25b* since the 1980s, with the frequency of these alleles exceeding ∼70% in the 2020s (Figure 4E, Supplementary Table S9).

#### 3.4.2 *Rht* haplotypes during pre– and post-Green Revolution periods

We observed 17 distinct *Rht* haplotypes based on 972 lines with complete genotype calls. Overall, the five most frequent haplotypes were *Rht*fZI*B1a/Rht*fZI*D1a/Rht24a/Rht25f* (n = 262), *Rht*fZI*B1b/Rht*fZI*D1a/Rht24a/Rht25f* (n = 184), *Rht*fZI*B1a/Rht*fZI*D1a/Rht24a/Rht25a* (n = 167), *Rht*fZI*B1a/Rht*fZI*D1b/Rht24a/Rht25f* (n = 106), and *Rht*fZI*B1a/Rht*fZI*D1a/Rht24b/Rht25b* (n = 60), accounting for roughly 75% of the panel (Supplementary Table S10). Before 1970, the two haplotypes, *Rht-B1a*/*Rht-D1a*/*Rht24a*/*Rht25a* (n = 137) and *Rht-B1a*/*Rht-D1a*/*Rht24a*/*Rht25f* (n = 136), were predominant in the region, while *Rht-B1b*/*Rht-D1a*/*Rht24a*/*Rht25f* (n = 181) was the most frequent after 1970 (Supplementary Table S11).

### 3.5 Effect of different *Rht* haplotypes on plant height

The plant height data from historical multi-environmental HRSWURN trials from 1968 to 2023 were used to study the effect of different *Rht* haplotypes. We had 653 genotypes with complete genotype calls and plant height BLUEs from multi-environment analysis to compare the effect of different *Rht* haplotypes on plant height. The distribution of plant height BLUEs is presented in Supplementary Figure S4. Based on pairwise comparisons, we observed substantial differences in plant height across different allelic combinations of four *Rht* genes (Figure 5). Lines carrying only *Rht-B1b* or *Rht-D1b* exhibited the highest individual effects on plant height, with mean heights of 80.6 cm and 79.7 cm, respectively, compared to 89.1 cm in lines with wild-type alleles at all four *Rht* loci. When these alleles were combined with dwarfing alleles at *Rht24b* and/or *Rht25b*, plant height was significantly reduced even further due to additive effects. For example, the shortest plants (75.4 cm) were observed when *Rht-D1b* was present in combination with *the Rht24b* and *Rht25b* alleles. Interestingly, the lines with wild-type alleles at *Rht-B1* and *Rht-D1* but having both *Rht24b* and *Rht25b* had a significant reduction in plant height (83.2 cm), compared to lines without dwarfing alleles (Figure 5).

**Figure 5.**
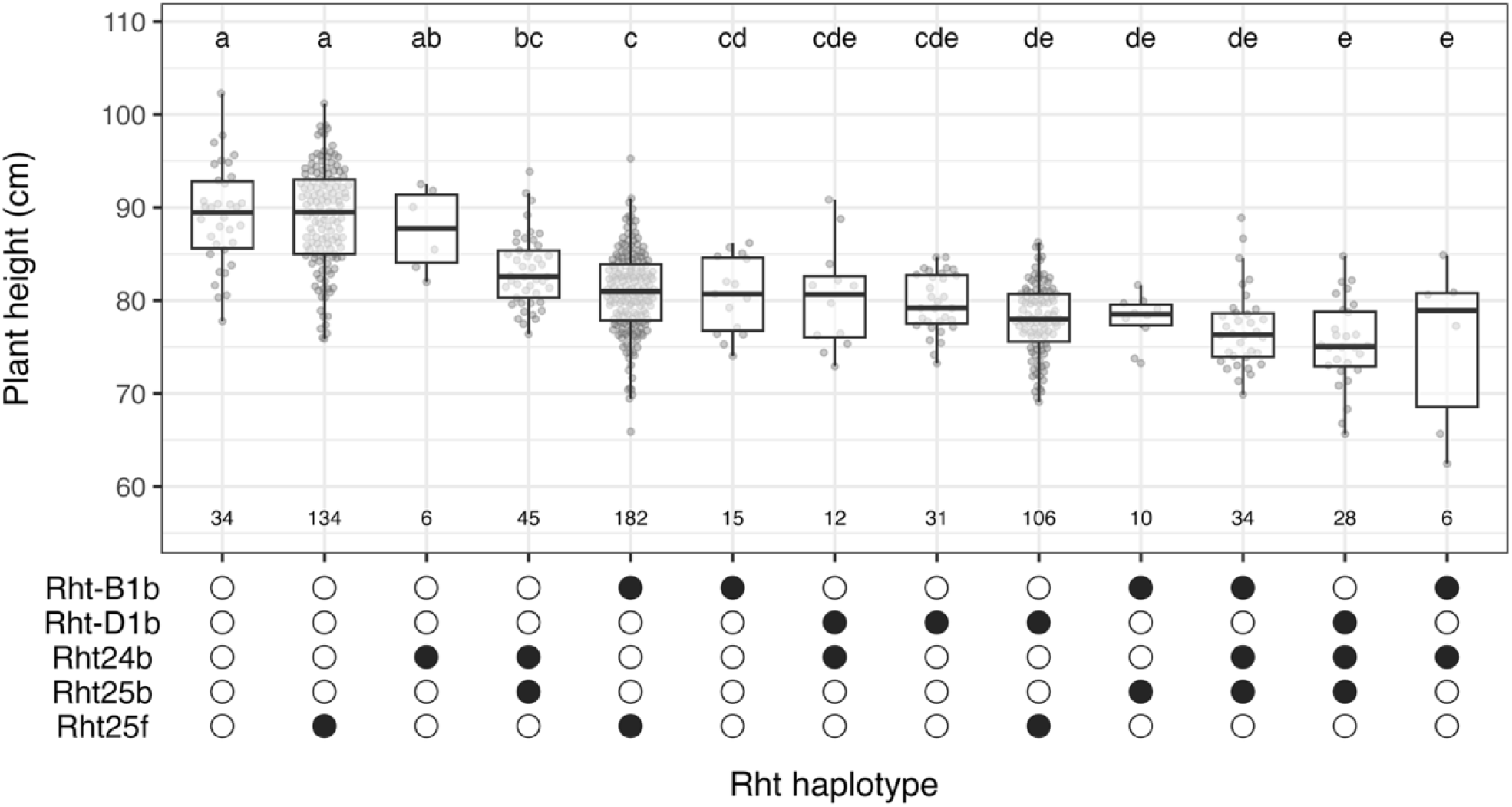
Effects of different *Rht* haplotypes (based on the *Rht-B1*, *Rht-D1*, *Rht24*, and *Rht25* genes) on plant height. The upper half overlays boxplots for plant height (BLUEs from multi-environment analysis) for every haplotype. The dot matrix in the lower panel indicates the allelic composition of each haplotype: a filled black circle denotes the presence of the listed dwarfing allele, while an open circle indicates its absence. Haplotypes are ordered from left to right by decreasing mean height. Compact-letter groups above each box come from a Tukey HSD test, and haplotypes sharing a letter do not differ significantly (α = 0.05). The numeral below every box is the number of lines in that haplotype class.

### 3.6 Allelic effects of different *Rht* loci in the North American HRSW region

Next, we used a linear gene-based model using plant height BLUEs from multi-environmental HRSWURN trials to statistically assess the substitution effects of different *Rht* genes and their interactions. Model selection was used to find the best linear model for plant height by comparing different combinations of the four *Rht* genes and their interactions (Supplementary Table S12). The best model, which included four *Rht* loci and three two-way interaction terms, explained up to 49% of the variability in plant height (Table 3). The intercept from the model reflects a mean plant height of 88.7 cm in the absence of dwarf alleles at the four *Rht* loci. The ANOVA from the linear model confirmed significant effects of *Rht-B1b* (–10.00 cm, p<0.001) and *Rht-D1b* (–10.65 cm, p<0.001), indicating these loci had the strongest dwarfing effect (∼12%). The alleles *Rht24b* (–2.70 cm, p<0.01) and *Rht25b* (–3.95 cm, p<0.001) also had a significant effect on plant height, although to a lesser extent than *Rht-B1* and *Rht-D1*. For *Rht25*, the *Rht25f* allele also showed a reduction in plant height, but the effect was statistically non-significant. The gene-based ANOVA results revealed significant genetic interactions, with *Rht-B1* showing a significant interaction with *Rht25*, and *Rht-D1* displaying a significant interaction with *Rht24* (Table 3). Among the two different *Rht25* alleles, *Rht25b* (p<0.05) showed stronger interaction with *Rht-B1b* compared to the *Rht25f* allele. Interestingly, the best models for individual locations (based on the highest *R^2^*) for all environments did not include the *Rht24b*:*Rht25b* interactions (Table 3), indicating a straightforward additive effect of combining dwarf alleles at this combination of genes.

**Table 3.** Fixed-effect estimates from the best-fit linear model describing the effect of four *Rht* loci on plant height (based on BLUEs obtained from multi-environment HRSWURN trials). The reference genotype for each locus is the tall, wild-type allele (*Rht-B1a*, *Rht-D1a*, *Rht24a*, *Rht25a*). Thus, the estimates represent the mean change in plant height (cm ± SE) associated with substituting the respective allele (or allele combination) for its reference, holding all other terms constant.

| **Genetic term** | **Estimate** | **Standard error** | **p-value** | **R<sup>2</sup>** |
| --- | --- | --- | --- | --- |
| Intercept | 89.87 | 0.59 | <0.001 | 0.49 |
| *Rht-B1b* | -10.00 | 1.16 | <0.001 |  |
| *Rht-D1b* | -10.65 | 0.52 | <0.001 |  |
| *Rht24b* | -2.70 | 0.83 | 0.001 |  |
| *Rht25b* | -3.95 | 0.93 | <0.001 |  |
| *Rht25f* | -1.06 | 0.61 | 0.083 |  |
| *Rht24b:Rht-D1b* | 3.12 | 1.05 | 0.003 |  |
| *Rht25b:Rht-B1b* | 3.33 | 1.45 | 0.022 |  |
| *Rht25f:Rht-B1b* | 2.06 | 1.23 | 0.094 |  |

### 3.7 Environment-specific allelic effects of different *Rht* loci

Furthermore, we used the best model from the previous analysis to estimate the allelic effects of *Rht* genes in four individual environments representing different moisture regimes within the North American HRSW region (Table 4). Among the four selected environments, STP had the highest average precipitation at 536 mm, followed by CRK at 351 mm, CRT at 334 mm, and WLT, which had the lowest at 262 mm. For comparison, we classified CRK, CRT, and WLT as dry environments due to their lower precipitation levels. The average plant height for tall genotypes (no dwarfing allele at four *Rht* genes) showed a trend with the annual precipitation, with the driest location, WLT, having the lowest plant height (Table 4). The ANOVA for individual environments revealed a consistent and significant effect of *Rht-B1b* and *Rht-D1b* across all environments with strong dwarfing effects. However, the effect of *Rht-B1b* and *Rht-D1b* was less pronounced in CRK, CRT, and WLT, with an average reduction of ∼9% in plant height, compared to ∼12% in the overall analysis (Table 4). The analysis also showed a significant effect of *Rht24b* in all the environments except for STP, suggesting the importance of this gene in dry regions of the Great Plains. The effect of *Rht25b* and *Rht25f* was environment-specific and typically smaller (Table 4). However, either one of the two *Rht25* dwarf alleles had a significant effect on plant height in dry environments (CRK, CRT, and WLT).

**Table 4.**
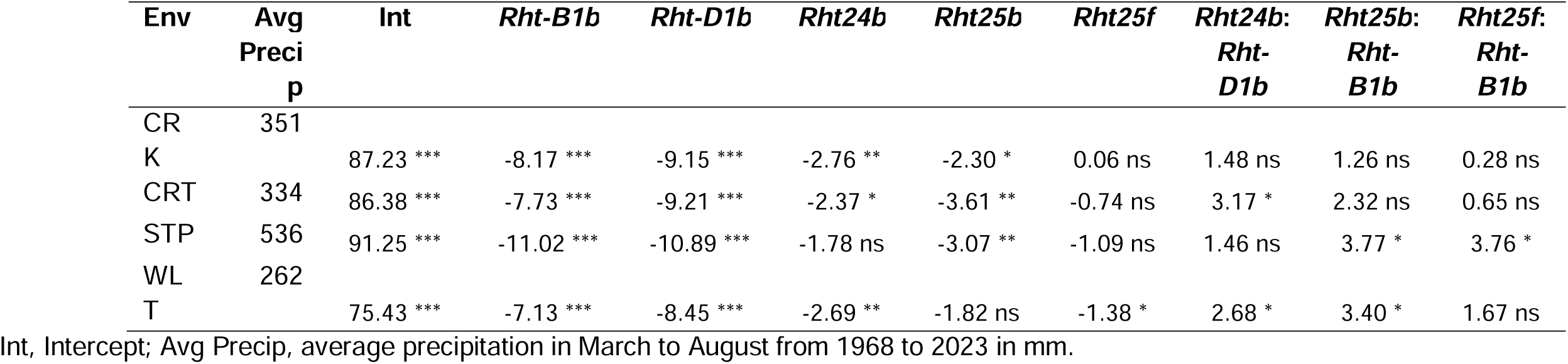
Allelic effects of gene-based terms included in the best-fit linear model for plant height across five diverse environments. The intercept indicates the mean plant height (cm) for each environment, excluding the effect of dwarfing alleles. The allelic effects listed for each dwarfing allele (*Rht-B1b*, *Rht-D1b*, *Rht24b*, *Rht25b*, and *Rht25f*) reflect their additive impact on plant height relative to the intercept. Interaction terms (e.g., *Rht24b*:*Rht-D1b*) denote additional non-additive effects observed when specific allele combinations are present. Statistical significance levels are indicated as follows: p < 0.05 (*), p < 0.01 (**), p < 0.001 (***); non-significant terms are marked as ns.

### 3.8 Genome-wide association analysis for plant height

A GWAS was performed to confirm the effect of major *Rht* genes and identify other putative genomic regions controlling plant height. A total of 11 marker-trait associations were identified based on a strict Bonferroni-corrected threshold (*p < 0.01*) that were located on seven different chromosomes (Figure 6; Supplementary Table S13). The two strongest associations were identified on chromosomes 4B (SNP chr4B_33614738, –log_10_(*P*) = 71.2) and 4D (SNP chr4D_19189840, –log_10_(*P*) = 39.3), respectively, corresponding to the *Rht-B1* and *Rht-D1* loci. The third significant association (SNP chr6A_431926237, –log_10_(*P*) = 12.4) was identified on chromosome 6A, which harbors the *Rht24* and *Rht25 loci*. In addition to these genes, we identified eight marker-trait associations on chromosomes 4A, 5A, 6B, and 7A (Figure 6; Supplementary Table S13), suggesting the role of other minor loci in controlling plant height.

**Figure 6.**
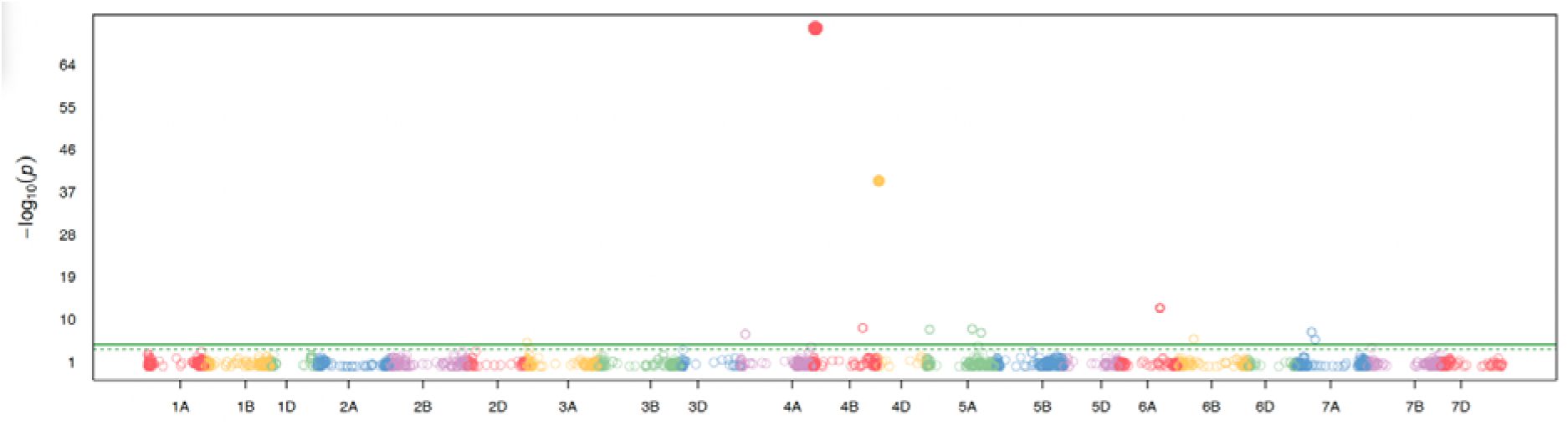
Manhattan plot presenting the marker-trait associations identified for plant height. The x-axis contains the 21 wheat chromosomes, and the y-axis represents the negative logarithm of p-values obtained from GWAS. The horizontal line elucidates the Bonferroni-corrected threshold of 5.36, while the dotted line elucidates the FDR-corrected threshold.

### 3.9 Allelic shifts in Vernalization (*Vrn*), photoperiod (*Ppd*), and other flowering-related genes

We observed temporal shifts in key genes associated with flowering time (Supplementary Figure S4; Supplementary Table S14). The frequency of the spring-type allele (*Vrn-A1a*) at the *Vrn-A1* locus has remained consistently high (>95%) since the 1930s, indicating early selection and fixation in spring wheat germplasm. Conversely, the winter-type alleles (*vrn-B1, vrn-D1*) at the *Vrn-B1* and *Vrn-D1* loci have historically dominated. The spring-type allele (*Vrn-D1a*) has shown a notable and consistent increase since the 1970s, reaching nearly 45% by the 2010s (Supplementary Figure S4; Supplementary Table S14). For *Vrn-D3*, the early-flowering allele (*Vrn-D3a*) was present in more than 60% of the germplasm until the 1950s, followed by a continuous decline to below 20% by the 2010s. In contrast, the allele associated with earliness at *FT3-B1* has been favored and was present in more than 90% of the germplasm from recent decades.

The *Ppd-D1a* allele that confers photoperiod insensitivity was introduced into the HRSW germplasm in the 1950s and increased to moderate frequencies (∼33%) by the 1990s (Figure 7; Supplementary Figure S4; Supplementary Table S14). However, the results indicate a selection toward the photoperiod-sensitive allele *Ppd-D1b* in the last two decades, with less than 20% lines from the 2020s carrying the insensitive allele. Interestingly, the deployment of *Ppd-D1a* was highly program-specific, with the MN program having this allele in about 75% of their germplasm in the 1990s and around 50% over the next two decades. In contrast, the MT and ND programs mainly relied on the photoperiod-sensitive *Ppd-D1b* allele (Supplementary Figure S5). Furthermore, another structural variant in the *Ppd-D1* gene (an MLE-insertion) has consistently increased across all breeding programs from the region, reflecting a preference for photoperiod sensitivity coupled with moderate earliness in recent decades (Supplementary Table S14). In addition to *Ppd-D1*, we found minimal variation in the major alleles at the *Ppd-A1* and *Ppd-B1* genes, which were excluded from this study.

**Figure 7.**
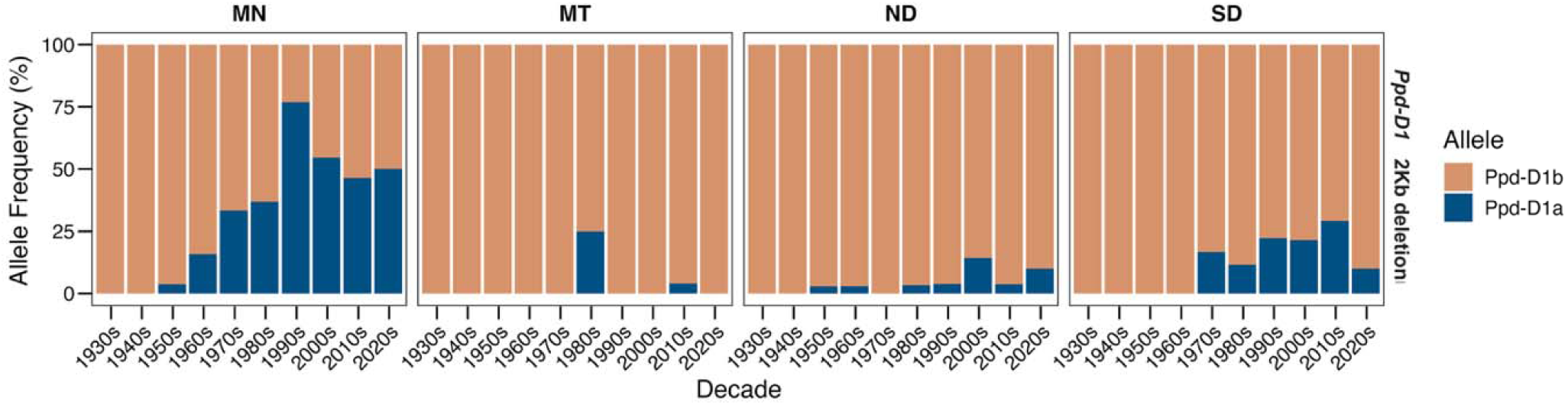
Temporal changes in the frequency of the 2 Kbp deletion variant (associated with photoperiod insensitivity) for the Ppd-D1 gene from the 1930s to the 2020s, separated by four major breeding programs: Minnesota, MN; Montana, MT; North Dakota, ND; and South Dakota, SD. Different alleles are distinguished by different colors, with blues representing alleles associated with photoperiod insensitivity and earliness.

## 4. Discussion

### 4.1 Plant breeding efforts mitigated early genetic diversity bottlenecks

HRSW breeding in the Northern Plains dates back to the late 19th century, when breeding efforts were initiated to address the challenges posed by the region’s harsh climate and disease pressures. The breeding efforts at the beginning of the 20th century relied on limited genetic diversity due to a very narrow foundational gene pool derived from a few ancestors (Mercado *et al*., 1996). Breeders successfully mitigated this bottleneck by introducing useful alleles from diverse sources. For example, adapted lines were crossed with introduced landraces like ‘Kota’ (CItr 5878) and ‘Webster’ (CItr 3780), hard red winter wheats like ‘Kanred’ (CItr 5146), and several interspecific crosses with tetraploid durum wheats like ‘Iumillo’. Notably, ‘Hope’ (CItr 13133) and ‘H-44’ were derived from crosses of Marquis with the ancient emmer ‘Yaroslav Emmer’ (McFadden, 1930; Klages, 1931). These early efforts resulted in a broadened genetic base and the development of popular cultivars such as ‘Thatcher’ (Hayes *et al*., 1936), which combined the adaptability of Marquis with the rust resistance of Kanred and Iumillo. Our results indicate that this genetic diversity was maintained in the 1930s and 1940s (Figure 3) when breeders heavily utilized Hope, H-44, Ceres, Marquis, Thatcher, and its sister line ‘Double Cross’ to develop new wheat cultivars.

Further, we observed a major increase in genetic variation during the 1950s and 1960s, attributed to independent events. In the late 1940s and early 1950s, breeders in the region introduced exotic wheats that provided resistance to stem and leaf rust and other valuable traits. Some of these introductions, such as ‘Frontana’ and ‘Surpresa’ from Brazil, or ‘Kenya Farmer’ and ‘Kenya 58’ from Kenya, were extensively used and led to the development of cultivars like ‘Conley’ (CItr 13157), ‘Justin’ (CItr 13462), ‘Polk’ (CItr 13773), and ‘Selkirk’ (CItr 13100). This was followed by the introduction of semi-dwarf wheats from CIMMYT, resulting in increased genetic diversity in 1950s and 1960s. At the same time, private seed companies may have helped maintain this genetic variation by introducing novel and exotic alleles into the HRSW region. The HRSWURN functioned largely under the wheat workers’ code of ethics until around 2000 (https://www.ars.usda.gov/ARSUserFiles/30421000/wheatcode.html), allowing breeding programs to cross with other entries in the nursery, which may have helped maintain regional genetic diversity. Finally, the FHB epidemics in the 1990s led breeders to introduce new sources of resistance that were often outside the traditional HRS germplasm, such as the Chinese wheat variety ‘Sumai 3’ (McMullen *et al*., 2012).

Earlier studies have explored genetic diversity in HRSW over time, but none of these studies utilized a large panel of lines from this class (Mercado *et al*., 1996; Fu *et al*., 2006; Sthapit *et al*., 2020, 2022; Semagn *et al*., 2021). Our results corroborate the findings that genetic diversity in the HRSW region increased in the late 1940s and this diversity has been maintained in subsequent decades. Our results do not show a sharp decline in genetic diversity in recent decades as reported by Sthapit et al. (2020) for HRSW from the Pacific Northwest. In the respective study, the authors used nine HRSW accessions from the Pacific Northwest, spanning the period from 2000 to 2019, to estimate genetic diversity, which may not accurately reflect the actual genetic diversity in the region over the given period.

We also observed some interesting patterns of relatedness among the major breeding programs. The MN program exhibited wide dispersion and maintained a broader genetic base compared to other programs, possibly due to the routine use of exotic sources for rust resistance and other traits. Further, the MN program showed strong relatedness with the PRIVATE germplasm owing to continuous germplasm exchange. The MN, ND, and SD public breeding programs were gifted germplasm from Pioneer Hi-Bred International, which closed its spring wheat program in the late 1980s (Busch *et al*., 1996), and our results data indicate that the MN program might have utilized this material at a higher frequency. The results also suggest a convergence among the major breeding programs over the past two decades, which may have an adverse impact on genetic variation in the years to come. Therefore, breeders must continue to monitor relatedness to prevent genetic erosion and maintain regional genetic diversity for sustained genetic gains.

### 4.2 FHB epidemics and adoption of Green Revolution genes in the Great Plains

The dwarfing alleles of two *Rht* homeologs, *Rht-B1b* and *Rht-D1b*, have been widely deployed in wheat breeding worldwide, although preferences have varied across different geographic and climatic regions (Guedira *et al*., 2010; Würschum *et al*., 2017). In the North American HRSW region, both alleles were first introduced in the 1960s (Figures 4A and 4B). It is noteworthy that none of the accessions from this study carried dwarfing alleles at both *Rht-B1* and *Rht-D1* loci. The *Rht-B1b* and *Rht-D1b* alleles are rarely found together in advanced material or released varieties because such combinations result in a “double dwarf” phenotype that is unacceptably short. Therefore, most of the double-dwarf lines resulting from crosses between parents segregating for both these loci are eliminated in the earlier generations. However, any alleles of *Rht24* or *Rht25* can be combined with *Rht-B1b* and *Rht-D1b* to produce a genotype with acceptable height.

After the introduction of dwarfing alleles at Rht-B1 and Rht-D1 in the 1960s, their frequencies increased with a general preference for *Rht-D1b* (Figures 4A and 4B). However, this trend reversed in the 1990s when most breeding programs shifted to *Rht-B1b*, and this transition period coincided with the devastating Fusarium head blight (FHB) epidemics in the Great Plains (McMullen *et al*., 2012). Recent studies have confirmed that *Rht-D1b* leads to significantly higher FHB severity than *Rht-B1b*, largely due to greater anther retention and increased Type I susceptibility (Buerstmayr and Buerstmayr, 2016, 2022). Although the association of *Rht-D1b* with FHB was not established at the time, our temporal analysis suggests that breeders selected against *Rht-D1b* and used the less detrimental *Rht-B1b*, illustrating how routine plant breeding influenced significant changes in allele frequencies for improved resistance to FHB.

In addition to regional adoption, we observed program-specific preferences for *Rht-B1* and *Rht-D1*, which may reflect adaptation to the diverse moisture regimes within the Great Plains region. The SD program maintained a low frequency of dwarf alleles at *Rht-B1* and *Rht-D1*, with only 20% of its post-1970 releases carrying the dwarf allele at either of these loci. Semi-dwarfing *Rht-B1b* and *Rht-D1b* alleles are known to perform best under optimal conditions but often incur a yield penalty in drought-prone environments (Worland and Snape, 2001; Würschum *et al*., 2017; Jatayev *et al*., 2020). Furthermore, our results indicate that the allelic effect of *Rht-B1b* and *Rht-D1b* on reducing plant height is diminished under dry conditions (Table 4). A previous study (Lanning *et al*., 2012*b*) also observed less height reduction due to *Rht-B1b* and *Rht-D1b* in environments with lower precipitation. It is likely that target environments for the SD program have not favored these semi-dwarf alleles, and breeders have strategically limited the use of *Rht-B1b* and *Rht-D1b*. Instead, they have incorporated alternative GA-sensitive dwarfing genes, such as *Rht24* and *Rht25*, which confer moderate height reduction without severely shortening coleoptiles (Grover *et al*., 2018; Jatayev *et al*., 2020; Pearce, 2021). These findings suggest that the already adapted *Rht24* and *Rht25* mutants can be effectively combined with other GA-sensitive dwarfing genes to breed for an optimal plant height for dry environments, without relying on the GA-insensitive *Rht* dwarfing genes.

### 4.3 Origin and distribution of *Rht24* and *Rht25* in the region

The temporal analysis of *Rht* genes in this study shows an early origin of *Rht24* and *Rht25* dwarfing alleles in the HRSW region of North America. The likely source of *Rht24b* in the region is the durum wheat introduction from Italy, Iumillo (PI 5996), which was crossed with Marquis in the 1920s to develop Marquillo (Supplementary Table S1), the first HRSW cultivar in the region to carry *Rht24b*. We acquired the seeds for Iumillo from the NPGS, and the KASP genotyping for *Rht24* confirmed that it carries the *Rht24b* allele (Supplementary Table S15). Nevertheless, our analysis suggests that this allele was mainly carried forward through ‘Double Cross’, a sister line of Thatcher that was extensively utilized in crossing blocks until the late 1940s.

According to Zhang et al. (2023), the most frequent *Rht25* mutation in a North American spring wheat collection was *Rht25b* (36.8%), followed by *Rht25f* (11.0%). In contrast, our results show a higher frequency of *Rht25f* in North American HRSW. Our results indicate that *Rht25f* came into this region through the founder cultivar Red Fife. The cultivar Marquis, a cross between Red Fife and ‘Hard Red Calcutta’ (Supplementary Table S1), carried the *Rht25f* allele. Hard Red Calcutta has been reported to carry a tall *Rht25a* allele (Zhang *et al*., 2023), which was also validated in this study, suggesting Red Fife may be the source. Another dwarf allele, *Rht25b*, was likely introduced to the HRSW region in the 1950s. All *Rht25b* positive lines during these years had the Brazilian introduction ‘Frontana’ (CItr 12470) in their pedigrees, with one notable exception, ‘N2292’ (Cltr 12967). We acquired the seed of Frontana and N2292 from NPGS, and both tested positive for *Rht25b*. Interestingly, none of the parents of N2292 (Pilot13/Merit//Lee) carried the *Rht25b* allele (Supplementary Table S15), and it is most likely that the seed of N2292 was contaminated. Hence, our results suggest that Frontana is the most likely source of *Rht25b* in the HRSW region, and the extensive use of this line as a parent, due to it containing the adult plant resistance gene *Lr34,* has contributed to an increase in the frequency of *Rht25b*. In agreement with Zhang et al. (2023), we also observed that *Rht25b* has increased gradually since the 1960s and replaced *Rht25f*.

### 4.4 Allelic effects of *Rht* genes and their genetic interactions

We observed that *Rht-D1b* had the strongest effect on reducing plant height, followed by *Rht-B1b*, with around a 12% reduction in plant height. However, the effect observed in our study is comparatively smaller than the previously reported ∼15-20% reduction in plant height by either of these genes (Jobson *et al*., 2019). Moreover, this effect is reduced further in dry environments. The *Rht24b* allele had a significant but mild effect on plant height compared to *Rht-B1b/Rht-D1b,* with a more pronounced effect in dry environments (Table 4). In contrast, *Rht25f*/b showed environment-specific effects, with *Rht25b* showing a significant effect on plant height compared to *Rht25f*. Zhang et al. (2023) have also observed that the effect of *Rht25f* on reducing plant height was non-significant. The allelic substitution analysis and positive selection for *Rht24b* and *Rht25b* in recent years suggest the promise of these GA-sensitive genes in the HRSW region of North America. For example, the average effect of *Rht-B1b* on plant height in three dry environments (CRK, CRT, and WLT) was 7.6 cm compared to 11 cm in STP. In the same three environments, the additive effect of dwarf alleles at *Rht24* and *Rht25* averaged around 5 cm (Table 4). Based on the trend between plant height and annual precipitation (Table 4), combining multiple GA-sensitive *Rht* genes could be a useful strategy to reduce plant height without incurring the adverse effects associated with GA-insensitive alleles at *Rht-B1* and *Rht-D1,* making it a suitable option for programs like SD.

The recent increase in *Rht24b* and *Rht25b* within backgrounds already carrying *Rht-B1b* or *Rht-D1b* remains unexplained. In this study, we observed significant genetic interactions between *Rht-B1* and *Rht25*, as well as between *Rht-D1* and *Rht24*. Zhang et at/ (2023) have also reported a significant interaction between *Rht25* and *Rht-B1*. It is possible that these interactions play a crucial role in optimizing plant height for various production environments, particularly the diverse moisture regimes of the Great Plains. For instance, our results suggest that combining *Rht-B1b* with *Rht24b* has an additive effect on reducing plant height, which could be useful in environments like STP. However, these are preliminary results based on historical data, and a comprehensive study is needed to investigate the impact of these genetic interactions on breeding for optimum plant height. Additionally, we propose a regional study using near-isogenic lines for these genes to enhance our understanding of the effects and interactions of *Rht* genes in adapted HRSW germplasm and explain the recent increase of these mutant alleles at *Rht24* and *Rht25*.

### 4.5 GWAS confirmed the role of *Rht* genes and suggested other loci controlling plant height

The analysis of historical data revealed substantial variation in plant height, particularly in lines lacking either the *Rht-B1* or *Rht-D1* genes (Figure 5). This motivated us to perform a GWAS to identify putative genomic regions controlling plant height in the HRSW growing region. The three strongest marker-trait associations were identified on chromosomes 4B, 4D, and 6A, which harbor *Rht-B1, Rht-D1, Rht24, and Rht25,* respectively. These results further confirm that these four loci play a crucial role in fine-tuning plant height in HRSW. We did not find any association on chromosomes 2D, which harbor *Rht8* (Gasperini *et al*., 2012), indicating that this gene is not significant in the HRSW region, and this is consistent with the genotyping of the *Rht8* marker in a subset of HRSW lines. In addition to the *Rht* genes, we identified several marker-trait associations on different chromosomes, suggesting the presence of other loci with minor effects that could be used to fine-tune plant height. It is important to acknowledge that the GWAS was conducted using a limited number of markers, and several regions may not have been adequately represented due to the limited genome coverage. However, the identification of known loci supports the validity of marker-trait associations, and these suggestive results could be used to confirm the identified regions in a further study.

### 4.6 Photoperiod sensitivity in high-latitude spring wheat

Despite a gradual increase in the frequency of the photoperiod-insensitive *Ppd-D1a* allele in the HRSW region until the 1990s, our results indicate a reversal of this trend over the last two decades. Moreover, the MT and ND public breeding programs have primarily relied on the photoperiod-sensitive *Ppd-D1b* allele. Earlier studies from the 1980s demonstrated that photoperiod-insensitive spring wheat lines yielded comparably to traditional photoperiod-sensitive lines, mainly due to their ability to initiate grain filling before the onset of extreme hot and dry spells in summer (Busch *et al*., 1984). Nevertheless, the photoperiod-sensitive lines have been shown to outyield their photoperiod-insensitive counterparts under early planting conditions, attributed to their increased vegetative growth period and improved biomass accumulation (Dyck *et al*., 2004; Lanning *et al*., 2012*a*). Given the recent trend toward earlier planting in the northern Great Plains, driven by warmer spring temperatures <u>(Lanning et al.</u> <u>2010)</u>, breeding programs appear to favor photoperiod-sensitive material to exploit longer vegetative phases and enhance yield potential. Additionally, the recent increases in the frequency of *Vrn-D1a*, the *Ppd-D1* MLE insertion allele, and other variants indicate that breeders are working to further fine-tune developmental timing for specific production environments.

### 4.7 Conclusion

By genotyping more than a thousand HRSWURN lines with a low-density 3K SNP array and key KASP markers, we have created a publicly accessible resource for wheat breeding programs. The genotype data enables the use of extensive publicly available data on agronomic and end-use quality traits collected from a large number of environments for genomic prediction and to dissect long-term genotype-by-environment-by-management interactions. Furthermore, the analysis of *Rht* genes in the Great Plains region provided valuable insights into the regional adoption of *Rht-B1b and Rht-D1b* alleles and supports the usefulness of deploying GA-sensitive alternatives, such as *Rht24* and *Rht25*, in semi-arid environments. Moreover, the genetic interactions among different *Rht* genes indicate that specific combinations can be successfully exploited to fine-tune plant height in various environments. For brevity, we primarily focused on the *Rht* genes and investigated their impact on plant height; however, the data will enable us to study other important genes, such as utilizing the long-term data on heading date to help us understand how flowering-time genetics have been adjusted in response to shifting agronomic regimes. In conclusion, this dataset offers a robust resource for both retrospective analysis of past breeding efforts and informing breeders in decision-making for improving the HRSW germplasm.

## Acknowledgment

The authors would like to thank the many researchers who have collaborated in the HRSWURN over the past century. We are also grateful to Dr. Jagdeep Sidhu from South Dakota State University for his input regarding methodology and the evolution of *Rht* genes.

## Author Contributions

HG, AR, and JA: conceptualization; HG and CB: methodology; HG and SB: data curation; HG: formal analysis; HG, SB, EC: investigation; JF, HG, SB, EC: genotyping and SNP calling; HG: visualization; HG: writing – original draft; SB, CB, EC, JF, KG, JC, AG, AR, JA: writing – review & editing; JA and AR: resources; JA: supervision; JA and AR: funding acquisition; and JA and AR: project administration.

## Disclaimer

The U.S. Department of Agriculture (USDA) prohibits discrimination in all its programs and activities on the basis of race, color, national origin, age, disability, and where applicable, sex, marital status, familial status, parental status, religion, sexual orientation, genetic information, political beliefs, reprisal, or because all or part of an individual’s income is derived from any public assistance program. Individuals with disabilities who require alternative means of communication for program information (such as Braille, large print, or audiotape) should contact USDA’s TARGET Center at (202) 720-2600 (voice and TDD). To file a complaint of discrimination, write to USDA, Director, Office of Civil Rights, 1400 Independence Avenue, S.W., Washington, D.C. 20250-9410, or call (800) 795-3272 (voice) or (202) 720-6382 (TDD). USDA is an equal opportunity provider and employer.

## Data Availability Statement

The datasets generated in this study are provided within the manuscript, its Supplementary files, or the repositories described in the paper. The genotype data, including the 3K array and KASP data, are available on the T3 database. The phenotype data for HRSWURN are available at the GrainGenes database (https://wheat.pw.usda.gov/GG3/germplasm). The corresponding authors should be contacted for any additional requests.

## Supplementary Data

### Tables

**Table S1.** Metadata for the 1,013 HRSWURN accessions.

**Table S2.** Description of functional markers associated with genes included in the USDA SoyWheOatBar 3K array

**Table S3.** KASP assays and genotype calls for key agronomic and quality loci.

**Table S4.** Geographic coordinates and annual precipitation for each of the four HRSWURN trial locations utilized in the plant height analysis.

**Table S5.** Distribution of high-quality SNPs across 21 wheat chromosomes.

**Table S6.** Summary of gene diversity (expected heterozygosity) estimated over the decades.

**Table S7.** Summary of gene diversity by breeding programs estimated over the decades.

**Table S8.** Temporal analysis of allele frequencies for *Rht* loci over the past century.

**Table S9.** Temporal analysis of allele frequencies for *Rht* loci over the past century, summarized by four major public breeding programs.

**Table S10.** Rht haplotype counts and frequencies across the entire panel.

**Table S11.** Rht haplotype frequencies in the pre– and post-1970 periods.

**Table S12.** Comparison of linear models for plant height (PH) using combinations of four dwarfing loci (*Rht-B1, Rht-D1, Rht24, Rht25*).

**Table S12.** Summary of the marker-trait associations identified for various end-use and processing traits.

**Table S13.** Allele frequencies for flowering time-related loci over the past century.

**Table S14.** KASP genotyping of selected historical lines for allelic discrimination of *Rht24* and *Rht25*.

### Figures

**Figure S1.** Density of 2,501 single-nucleotide polymorphism (SNP) markers across the 21 wheat chromosomes.

**Figure S2.** Bayesian STRUCTURE barlZIplot (KL=L2) for 1,013 HRSW accessions based on 3K SNPs filtered for <5% MAF (n = 2,297 SNPs).

**Figure S3.** Phenotypic distribution of plant height (PH) BLUEs from multi-environment analysis in 653 genotypes with complete genotype calls.

**Figure S4.** Changes in allele frequency of selected genes related to flowering time.

**Figure S5.** Temporal changes in the frequency of three different structural variants for the *Ppd-D1* gene from the 1930s to 2020s, separated by four major breeding programs, including Minnesota, MN; Montana, MT; North Dakota, ND; South Dakota, SD.

