## Supplemental_Figures for "A century of breeding has preserved genetic variation, accumulated favorable alleles, and shaped the *Rht* genes portfolio in North American spring wheat"

### Supplementary Figures

**Figure S1.** Density of 2,501 single-nucleotide polymorphism (SNP) markers across the 21 wheat chromosomes.

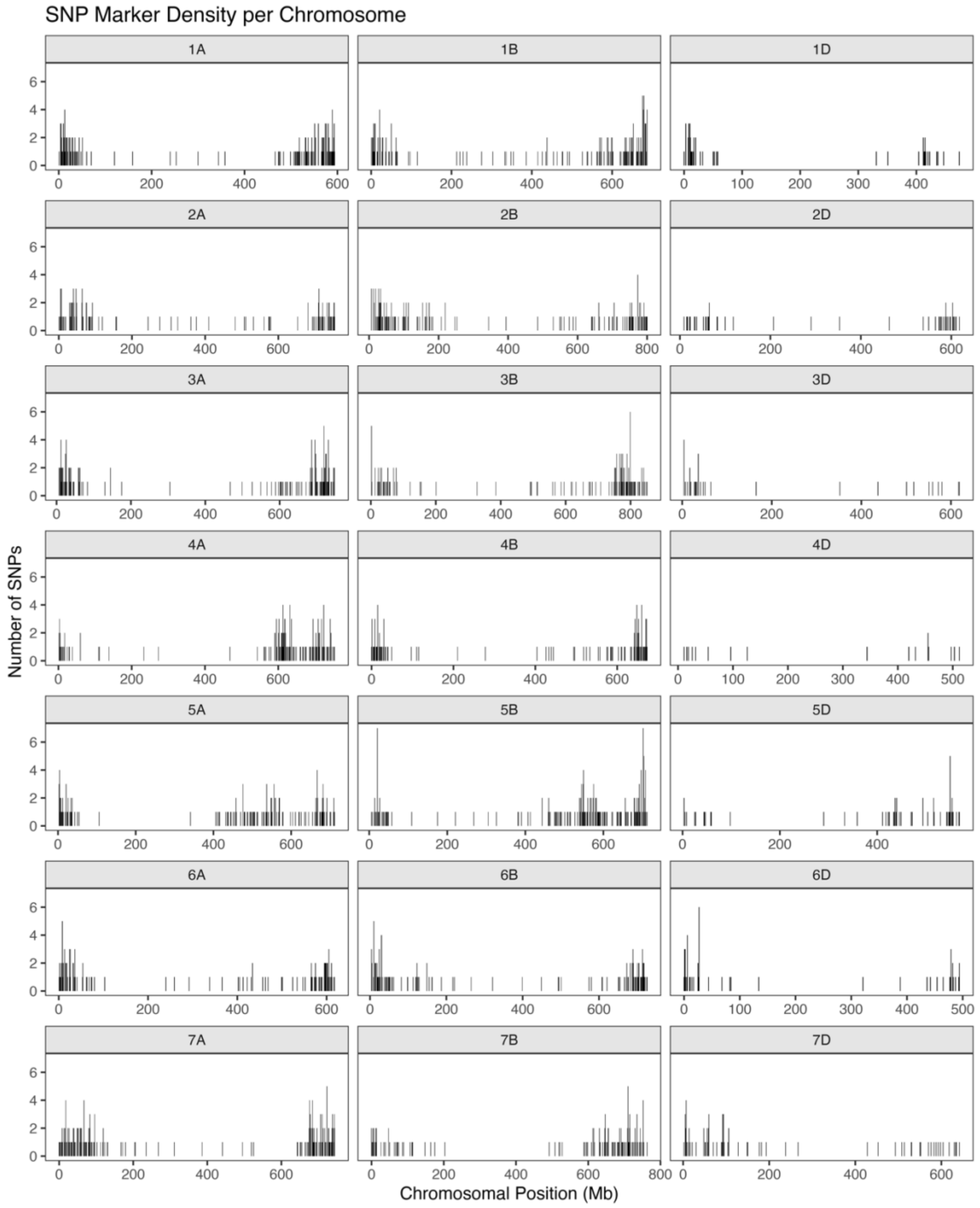

**Figure S2.** Bayesian STRUCTURE bar-plot (K = 2) for 1,140 HRS wheat accessions based on 3K SNPs filtered for <5% MAF (n = 2,297 SNPs). Each vertical bar represents one genotype, partitioned according to its estimated membership coefficient (Q) in the two inferred genetic clusters (Q1, light-blue; Q2, orange). Accessions are ordered by increasing proportion of Q2 ancestry. The horizontal strip below the STRUCTURE plot classifies the same individuals as historical (Pre-1970, blue) or modern (Post-1970, green) entries.

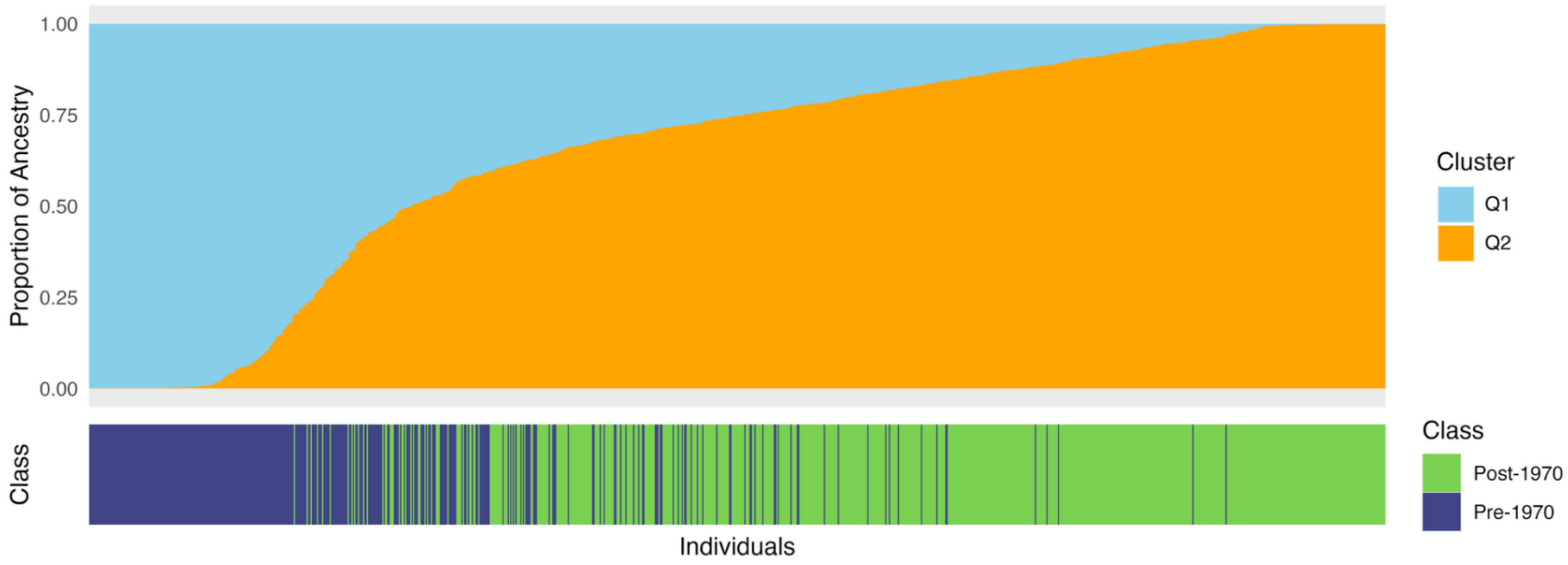

**Figure S3.** Phenotypic distribution of plant height (PH, centimeters) BLUEs from multi-environment analysis in 653 genotypes with complete genotype calls. a) Individual PH observations plotted by sample index (Individual on the x-axis and observed PH on the y-axis. (b) Box-and-whisker plot summarizing PH. (c) Histogram of PH showing observation frequencies. (d) Kernel density estimate of the PH distribution. (e) Empirical cumulative distribution function (ECDF) of PH.

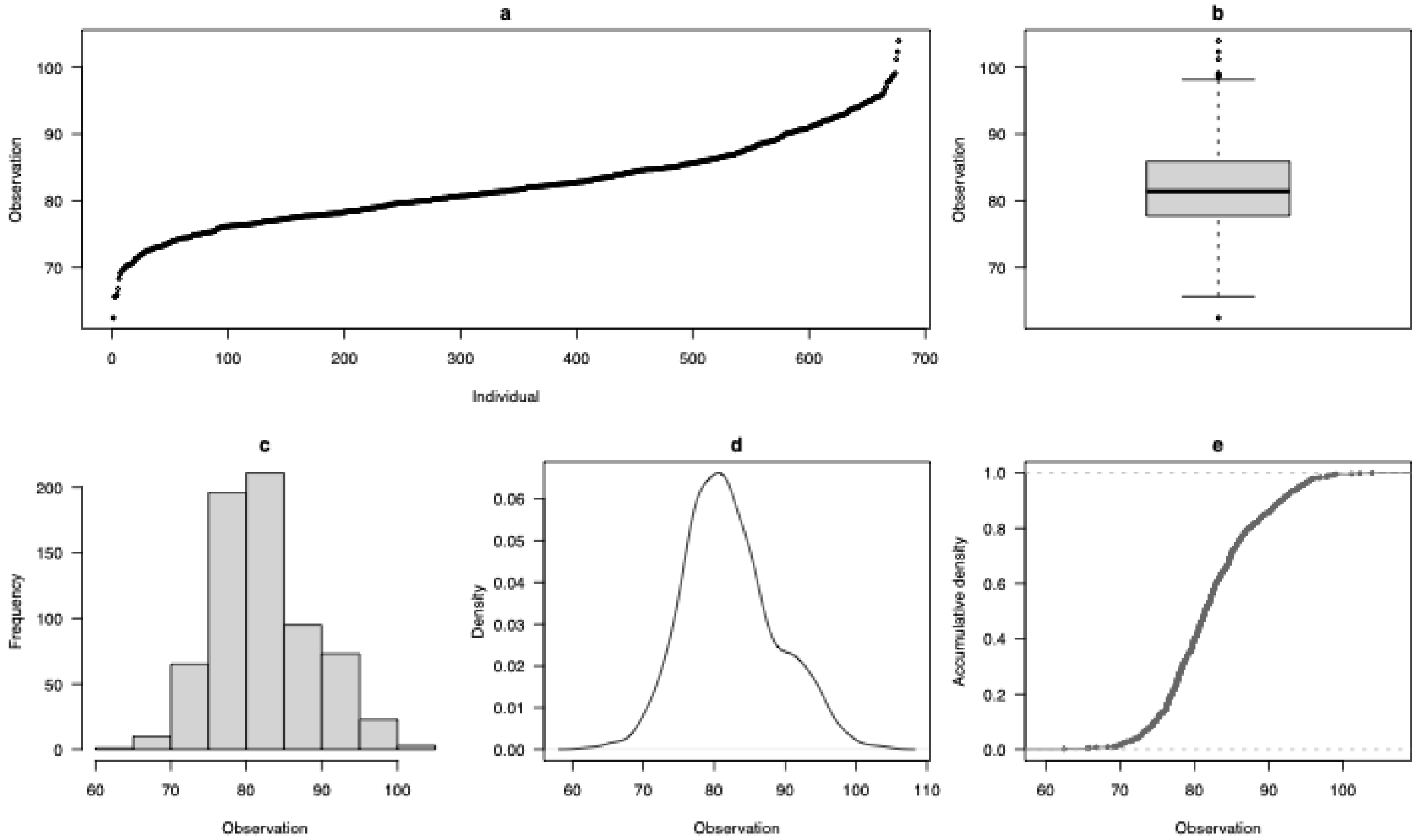

**Figure S4.** Changes in allele frequency of important genes related to flowering time across decades. Panels show frequency changes for various allelic variants in Ppd-D1 (presence or absence of either of 2KB deletion, MLE insertion, Norstar deletion), FT3-B1 (alleles associated with late or early flowering), and Vrn-A1, Vrn-B1, Vrn-D1, and Vrn-D3 ('a' allele at each Vrn loci is associated with spring-type or early flowering). Different alleles at each locus are represented by different colors, with blues representing alleles associated with earliness.

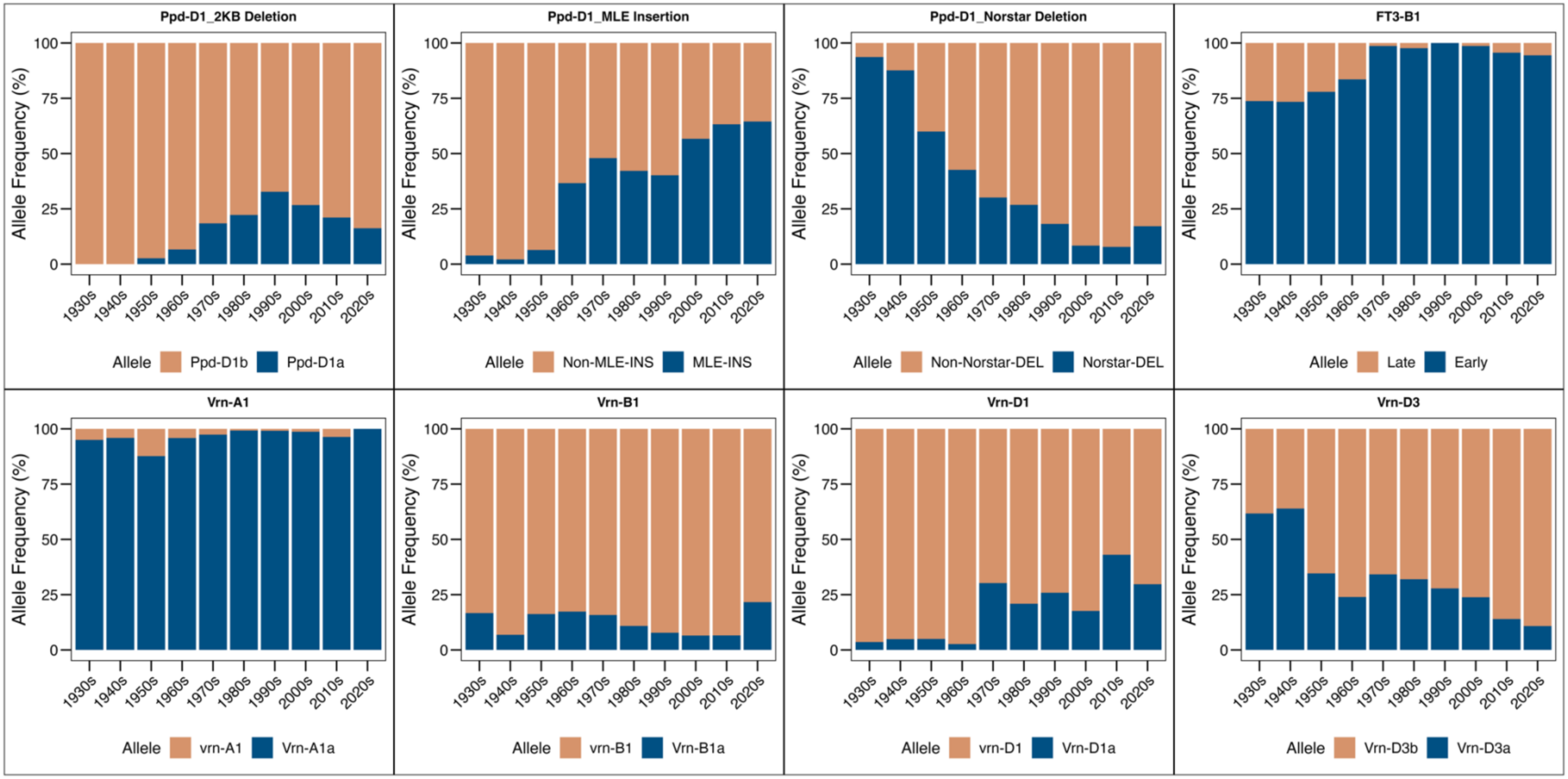

**Figure S5.** Temporal changes in the frequency of three different structural variants for the Ppd-D1 gene from the 1930s to 2020s, separated by four major breeding programs, including Minnesota, MN; Montana, MT; North Dakota, ND; South Dakota, SD. Panels display changes for specific allelic variants: Ppd-D1 2KB deletion (variant associated with photoperiod sensitivity), Ppd-D1 mariner-like transposable element (MLE) insertion in intron 1, and Ppd-D1 5-base pair deletion in exon 7 (Norstar deletion). Different alleles are distinguished by different colors, with blues representing alleles associated with earliness.

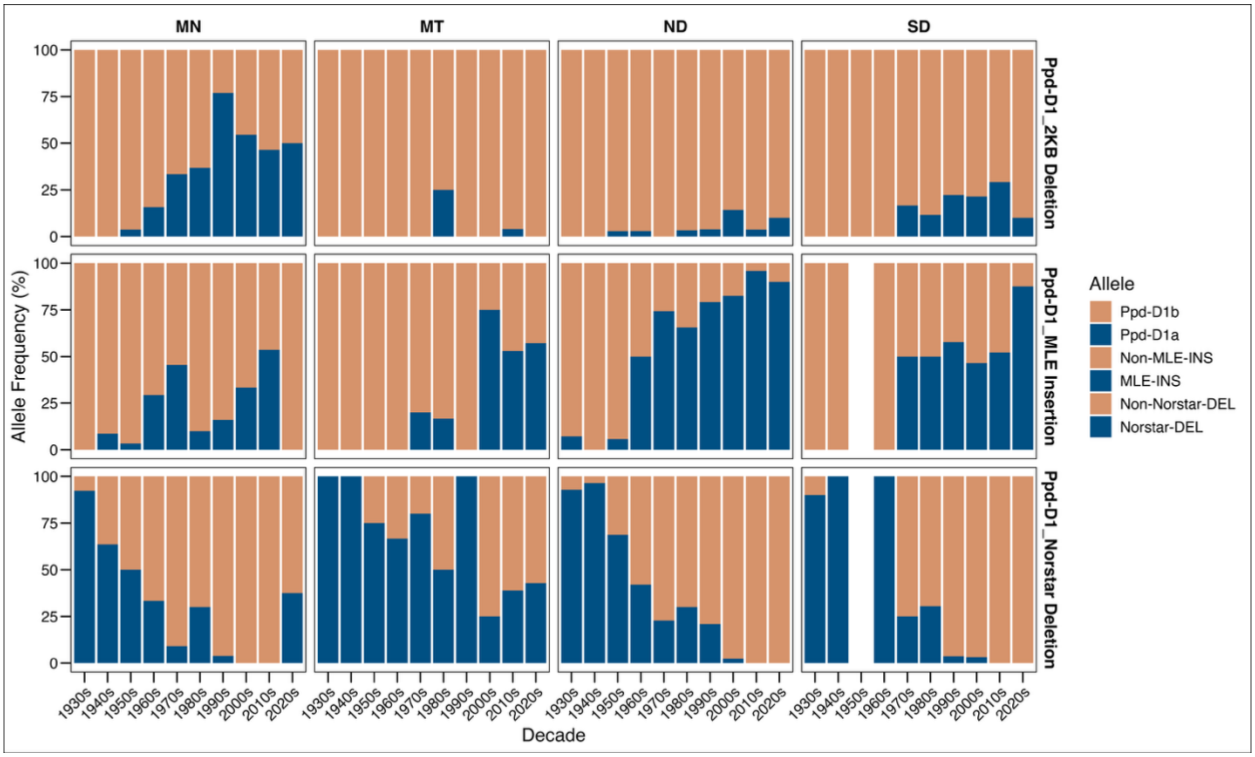
